# Brain Aging in Major Depressive Disorder: Results from the ENIGMA Major Depressive Disorder working group

**DOI:** 10.1101/560623

**Authors:** Laura K M Han, Richard Dinga, Tim Hahn, Christopher R K Ching, Lisa T Eyler, Lyubomir Aftanas, Moji Aghajani, André Aleman, Bernhard T Baune, Klaus Berger, Ivan Brak, Geraldo Busatto Filho, Angela Carballedo, Colm G Connolly, Baptiste Couvy-Duchesne, Kathryn Cullen, Udo Dannlowski, Christopher G Davey, Danai Dima, Fabio L S Duran, Verena Enneking, Elena Filimonova, Stefan Frenzel, Thomas Frodl, Cynthia H Y Fu, Beata R Godlewska, Ian H Gotlib, Hans J Grabe, Nynke A Groenewold, Dominik Grotegerd, Oliver Gruber, Geoffrey B Hall, Ben J Harrison, Sean N Hatton, Marco Hermesdorf, Ian B Hickie, Tiffany C Ho, Norbert Hosten, Andreas Jansen, Claas Kähler, Tilo Kircher, Bonnie Klimes-Dougan, Bernd Krämer, Axel Krug, Jim Lagopoulos, Ramona Leenings, Frank P MacMaster, Glenda MacQueen, Andrew McIntosh, Quinn McLellan, Katie L McMahon, Sarah E Medland, Bryon A Mueller, Benson Mwangi, Evgeny Osipov, Maria J Portella, Elena Pozzi, Liesbeth Reneman, Jonathan Repple, Pedro G P Rosa, Matthew D Sacchet, Philipp G Sämann, Knut Schnell, Anouk Schrantee, Egle Simulionyte, Jair C Soares, Jens Sommer, Dan J Stein, Olaf Steinsträter, Lachlan T Strike, Sophia I Thomopoulos, Marie-José van Tol, Ilya M Veer, Robert R J M Vermeiren, Henrik Walter, Nic J A van der Wee, Steven J A van der Werff, Heather Whalley, Nils R Winter, Katharina Wittfeld, Margaret J Wright, Mon-Ju Wu, Henry Völzke, Tony T Yang, Vasileios Zannias, Greig I de Zubicaray, Giovana B Zunta-Soares, Christoph Abé, Martin Alda, Ole A Andreassen, Erlend Bøen, Caterina M Bonnin, Erick J Canales-Rodriguez, Dara Cannon, Xavier Caseras, Tiffany M Chaim-Avancini, Torbjørn Elvsåshagen, Pauline Favre, Sonya F Foley, Janice M Fullerton, Jose M Goikolea, Bartholomeus C M Haarman, Tomas Hajek, Chantal Henry, Josselin Houenou, Fleur M Howells, Martin Ingvar, Rayus Kuplicki, Beny Lafer, Mikael Landén, Rodrigo Machado-Vieira, Ulrik F Malt, Colm McDonald, Philip B Mitchell, Leila Nabulsi, Maria Concepcion Garcia Otaduy, Bronwyn J Overs, Mircea Polosan, Edith Pomarol-Clotet, Joaquim Radua, Maria M Rive, Gloria Roberts, Henricus G Ruhe, Raymond Salvador, Salvador Sarró, Theodore D Satterthwaite, Jonathan Savitz, Aart H Schene, Peter R Schofield, Mauricio H Serpa, Kang Sim, Marcio Gerhardt Soeiro-de-Souza, Ashley N Sutherland, Henk S Temmingh, Garrett M Timmons, Anne Uhlmann, Eduard Vieta, Daniel H Wolf, Marcus V Zanetti, Neda Jahanshad, Paul M Thompson, Dick J Veltman, Brenda W J H Penninx, Andre F Marquand, James H Cole, Lianne Schmaal

## Abstract

**Background:** Major depressive disorder (MDD) is associated with an increased risk of brain atrophy, aging-related diseases, and mortality. We examined potential advanced brain aging in MDD patients, and whether this process is associated with clinical characteristics in a large multi-center international dataset.

**Methods:** We performed a mega-analysis by pooling brain measures derived from T1-weighted MRI scans from 29 samples worldwide. Normative brain aging was estimated by predicting chronological age (10-75 years) from 7 subcortical volumes, 34 cortical thickness and 34 surface area, lateral ventricles and total intracranial volume measures separately in 1,147 male and 1,386 female controls from the ENIGMA MDD working group. The learned model parameters were applied to 1,089 male controls and 1,167 depressed males, and 1,326 female controls and 2,044 depressed females to obtain independent unbiased brain-based age predictions. The difference between predicted “brain age” and chronological age was calculated to indicate brain predicted age difference (brain-PAD).

**Findings:** On average, MDD patients showed a higher brain-PAD of +0.90 (SE 0.21) years (Cohen’s d=0.12, 95% CI 0.06-0.17) compared to controls. Relative to controls, first-episode and currently depressed patients showed higher brain-PAD (+1.2 [0.3] years), and the largest effect was observed in those with late-onset depression (+1.7 [0.7] years). In addition, higher brain-PAD was associated with higher self-reported depressive symptomatology (b=0.05, p=0.004).

**Interpretation:** This highly powered collaborative effort showed subtle patterns of abnormal structural brain aging in MDD. Substantial within-group variance and overlap between groups were observed. Longitudinal studies of MDD and somatic health outcomes are needed to further assess the predictive value of these brain-PAD estimates.

**Funding:** This work was supported, in part, by NIH grants U54 EB020403 and R01 MH116147.

## Research in context

### Evidence before this study

Accumulating evidence from studies suggests that, at the group level, MDD patients follow advanced aging trajectories, as their functional (e.g. walking speed, hand grip strength) and biological state (e.g. telomeres, epigenetics, mitochondria) reflects what is normally expected at an older age (i.e. biological age “outpaces” chronological age). While subtle structural brain abnormalities have been identified in MDD, it remains to be elucidated whether patients also deviate from the normal aging process at the brain level (brain predicted age difference [brain-PAD]) and whether this deviation is associated with clinical characteristics. We searched PubMed for relevant literature published in English [Language] before January 25, 2019. In this search we used ((‘brain age’ OR ‘brainAGE’ OR ‘brain-PAD’ OR ‘predicted brain ag*’) AND ‘depression’ [Title/Abstract]), which revealed only two papers. One study found that MDD patients (N=104) were estimated to be +4.0 years older using brain-based age prediction models. A second study reported a non-significant relationship between brain-PAD and a short self-report scale of depressive symptoms in male veterans (N=359) who served in the United States military. Thus, whether a diagnosis of MDD is associated with the multivariate metric of brain aging in a large dataset, and which clinical characteristics further impact this metric, remains elusive.

### Added value of this study

To our knowledge, this is the first study to examine deviations of normative brain aging in MDD and associated clinical heterogeneity in a large international and multi-center dataset, by pooling data from >8,000 subjects from 29 research samples worldwide. The current study shows that chronological age can be predicted from gray matter features in a large heterogeneous dataset with an age range covering almost the entire lifespan (10-75 years). Moreover, we show that our brain age prediction model generalizes to unseen hold-out samples, as well as to completely independent samples from different scanning sites. We found that, at the group level, patients had, on average, a +0.90 years greater discrepancy between their predicted and actual age compared to control participants and there was a subtle relationship between self-reported symptom severity and advanced brain aging in the MDD group. Finally, the strongest effects were observed in patients with a late onset of depression (>55 years old; +1.7 years), currently depressed (+1.2 years), and in their first episode (+1.2 years), compared to controls.

### Implications of all the available evidence

This study confirms previously observed advanced biological aging in MDD at the group and brain level of analysis. However, it is important to mention the large within-group and small between-group variance, demonstrating that many patients did not show advanced brain aging. Our work contributes to the maturation of brain age models in terms of generalizability, deployability, and shareability, in pursuance of a canonical brain age algorithm. Further, other research groups with deep clinical phenotyping and longitudinal information on mental and somatic health outcomes may use our model to promote continued growth of knowledge for greater clinical application.

## Introduction

Major Depressive Disorder (MDD) is associated with an increased risk of cognitive decline,^1^ brain atrophy,^2^ aging-related diseases,^2^ and importantly, overall mortality.^3,4^ While normal aging is associated with significant loss of gray matter,^5^ growing evidence suggests that neuropsychiatric disorders such as depression may have an accelerating effect on age-related brain atrophy.^6^ Simultaneously, the aging population is increasing, and both depression and aging have been linked to poor somatic health and quality of life, and increased costs for society and healthcare.^7,8^ This underscores the importance of identifying brain aging patterns in MDD patients to determine whether and how they deviate from healthy patterns of aging.

Emerging evidence indicates that chronological age and biological age may be distinct processes that can diverge. Current multivariate pattern methods can predict chronological age from biological data (i.e., epigenetics, transcriptomics, proteomics, metabolomics, see Jylhava, Pedersen, and Hagg for a review)^9^ with high accuracy. Similarly, chronological age can be predicted from brain images, resulting in an estimate known as “brain age”.^10^ Importantly, by calculating the difference between a person’s estimated brain age and their chronological age, one can translate a complex aging pattern across the brain into a single outcome:^11^ brain-predicted age difference (brain-PAD).^12^ A positive brain-PAD represents having an ‘older’ brain than expected for a person of their chronological age, whereas a negative brain-PAD signals a ‘younger’ brain than expected at the given chronological age. Higher brain-PAD scores have been associated with greater cognitive impairment,^13^ increased morbidity,^10^ and exposure to cumulative negative fateful life events (e.g., death of a close family member, financial hardship, or divorce).^14^

Prior studies from the Enhancing NeuroImaging Genetics through Meta-analysis (ENIGMA)-MDD consortium with sample sizes over 9,000 participants have shown subtle reductions in subcortical structure volumes in major depression that were robustly detected across many samples worldwide. Specifically, smaller hippocampal volumes were found in individuals with earlier age of onset and recurrent episode status.^15^ In addition, different patterns of cortical alterations were found in adolescents versus adults with MDD, suggesting that MDD may affect brain morphology (or vice versa) in a way that depends on the developmental stage of the individual.^16^ Likewise, brain development and aging likely differ by sex.^17^ The different neural and clinical presentations of depression and aging across sex emphasize the need to stratify populations studied into groups of females and males to better understand sex-dependent or sex-specific effects.

Given that prior studies suggest advanced biological aging in MDD (e.g., shorter telomere length,^18^ greater epigenetic aging,^19,20^ and advanced brain aging),^6^ it is important to examine whether biological aging findings in depression can be confirmed in a large heterogeneous dataset consisting of many independent samples worldwide, based on commonly derived gray matter measures. Only a handful of studies have investigated brain-PAD in people with psychiatric disorders,^21^ showing older brain-PAD in schizophrenia,^6,22,23^ borderline personality disorder, and first-episode and at-risk mental state for psychosis,^6,24^ yet findings were less consistent in bipolar disorder.^23,25^

Only two studies to date specifically investigated premature brain aging in MDD - using relatively small samples of 104 and 211 patients, respectively, with inconsistent findings of a brain-PAD of +4.0 years versus no significant difference.^6,26^ The current study is the first to examine brain aging in over 8,000 individuals from the ENIGMA MDD consortium (29 cohorts, 11 countries worldwide), covering almost the entire lifespan (10-75 years). We hypothesized higher brain-PAD in MDD patients compared to controls. We also conducted exploratory analyses to investigate whether higher brain-PAD in MDD patients was associated with demographic (age, sex) and clinical characteristics such as disease recurrence, antidepressant use, remission status, depression severity, and age of onset of depression.

## Methods

### Samples

Twenty-nine cohorts from the ENIGMA-MDD working group with neuroimaging and clinical data from MDD patients and controls participated in this study (**appendix**). The combined sample covered almost the entire lifespan (10-75 years of age). Details regarding demographics, clinical characteristics, and exclusion criteria for each cohort may be found in the **appendix**. Because the literature suggests differential brain development and maturation by sex,^17^ we estimated brain age models separately for male and female samples. Sites with less than ten healthy males or females were excluded from the training dataset and subsequent analyses (for exclusions see **appendix**). In total, we included data from N=8,159 (93.5%) participants, including N=4,948 (56.7%) control individuals (N=2,236 [45.2%] males; N=2,712 [54.8%] females) and N=3,211 (36.8%) individuals with MDD (N=1,167 [36.3%] males; N=2,044 [63.7%] females). All participating sites obtained approval from the appropriate local institutional review boards and ethics committees, and all study participants or their parents/guardians provided written informed consent.

### Training and test samples

An overview of the data partition is shown in **figure 1A** and described in more detail in the **appendix**. Structural brain measures from 1,147 male obtained from 28 scanners and 1,386 female controls obtained from 34 scanners were included in the training sample. The top panel in **figure 1B** shows the chronological age distribution in the training sample. A hold-out dataset comprised of controls served as test sample to validate the accuracy of brain age prediction model; 1,089 male and 1,326 female controls from the same scanning sites were included. Likewise, 1,167 male and 2,044 female MDD patients from the corresponding neuroimaging sites were included in the MDD test sample. The bottom panel in **figure 1B** shows the chronological age distributions across the test samples. More details on data partitioning are shown in the **appendix**.

**Figure 1:**
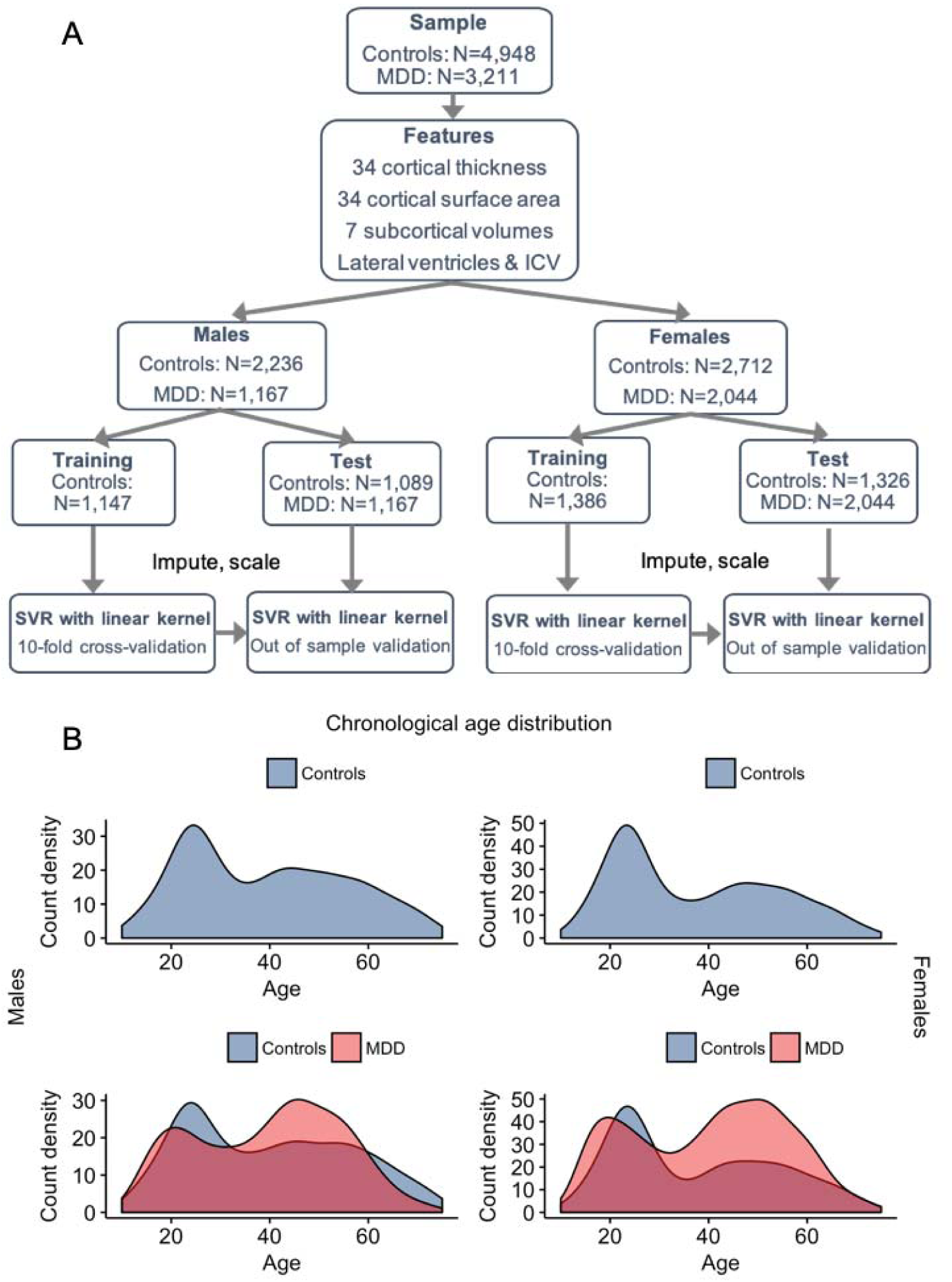
(A) Schematic illustration of features used and data partition into training and test samples, separately for males and females. **(B)** Data from control groups (*blue*) were partitioned within scanning sites preserving chronological age distribution. Major depressive disorder (MDD) groups are shown in *red*. The *top panel* illustrates the male and female training samples. The *bottom panels* show the male (controls: mean [SD] in years, 40.0 [16.5]; MDD: 39.6 [14.8]) and female test samples (controls: 37.6 [16.2]; MDD: 40.0 [15.5]). ICV, intracranial volume; SVR, support vector regression.

### Image processing and analysis

Structural T1-weighted scans of each subject were acquired at each site and analyzed locally using standardized protocols to facilitate harmonized image analysis across multiple sites (http://enigma.ini.usc.edu/protocols/imaging-protocols/). Briefly, the fully automated and validated segmentation software, FreeSurfer 5.1 or 5.3 was used to segment seven subcortical gray matter regions (nucleus accumbens, amygdala, caudate, hippocampus, pallidum, putamen, and thalamus), lateral ventricles, 34 cortical thickness and 34 surface area measures, and total intracranial volume (ICV). Segmentations were visually inspected and statistically examined for outliers. Further details on cohort type, image acquisition parameters, software descriptions, and quality control may be found in the **appendix**. Individual-level structural brain measures and clinical and demographic measures from each cohort were pooled at a central site to perform the mega-analysis.

### Brain age prediction model

To estimate the normative brain age models, we combined the FreeSurfer measures from the left and right hemispheres by calculating the mean ((left+right)/2) of volumes for subcortical regions and lateral ventricles, and thickness and surface area for cortical regions. Using a mega-analytic approach, we first estimated normative models of the association between the 77 average structural brain measures and chronological age in the training sample of controls (separately for males and females) using a support vector regression (SVR) with a linear kernel, from the python-based *sklearn* package.^27^ All measures were combined as predictors in a single multivariate model.

To assess model performance and optimize the regularization parameter, C, we performed 10-fold cross-validation. To quantify model performance, we calculated the mean absolute error (MAE) between predicted brain age and chronological age. Both male and female brain age models will be made public upon publication (https://www.photon-ai.com/); for guidelines and instructions, see **appendix**. Of note, we also estimated a model including left and right hemisphere measures, that did not result in significantly superior prediction accuracy, which allowed us to reduce the feature space to average left/right values as described (data not shown). We also compared the SVR to other machine learning methods, including ridge regression, Gaussian process regression, and generalized additive models. Results of these comparisons are provided in the **appendix**; briefly, the different approaches all showed similar performance to the model presented here.

### Model validation

Model performance was further validated in the test sample of controls. The parameters learned from the trained model in controls were applied to the test sample of controls and to the MDD test samples to obtain brain-based age estimates for these individuals. To assess model performance in these test samples, we calculated: a) MAE; b) Pearson correlation coefficients between predicted brain age and chronological age; and c) the proportion of the variance explained by the model (R²). To evaluate generalization power to completely independent test samples, we also applied the training model parameters to healthy control subjects (males, N=646; females, N=757) from the ENIGMA Bipolar Disorder (BD) working group (**appendix)**.

### Statistical analyses

All statistical analyses were conducted in the test samples only. Brain-PAD (predicted brain-based age - chronological age) was calculated for each individual and used as the outcome variable. While different prediction models were built for males and females separately, the generated brain-PAD estimates were pooled for statistical analyses. For our main analysis, we investigated three linear mixed models (LMM) of brain-PAD: a) main effects of age, sex, and diagnosis, b) all main effects and all second order interactions of age, sex, and diagnosis, and c) main effects and all second and third order interactions of age, sex, and diagnosis. To calculate the association between each FreeSurfer feature and brain-PAD, we used univariate regressions corrected for multiple comparisons (false discovery rate; FDR). Surface area and subcortical measures were additionally corrected for ICV.

Within MDD patients, we also used LMM to examine associations of brain-PAD with clinical characteristics, including recurrence status (first vs. recurrent episode), antidepressant use at time of scanning (yes/no), remission status (currently depressed vs. remitted), depression severity at study inclusion (the 17-item Hamilton Depression Rating Scale (HDRS-17) and the Beck Depression Inventory (BDI-II)), and age of onset of depression (categorized as: early, <26 years; adult, >25 & <56 years; and late onset, >55 years). All analyses included scanning site as a random intercept to account for scanner and FreeSurfer version differences and were corrected for chronological age, age^2^, age^3^, and sex, tested two-sided. Findings were considered statistically significant at *p*<0.05.

### Role of the funding source

The study design, data collection, analysis, interpretation, writing, and submission of this report were performed independently from any funding source. The corresponding author had full access to the complete dataset in the study. All authors had the final responsibility for the decision to submit for publication.

## Results

### Brain age can be predicted from regional brain measures

Within the training set of controls, under cross-validation the structural brain measures predicted chronological age with a MAE of 6.86 (SD 5.32) years in males and 6.91 (5.34) years in females. Correlations between chronological and predicted brain age were r=0.85, p<0.001 in males, and r=0.84, p<0.001 in females, with R^2^=0.72 and R^2^=0.71, respectively. When applying the model parameters to the test samples of controls, the MAE was 6.35 (4.92) and 6.63 (5.08) years for males and females, respectively. Similarly, within the MDD group, the MAE was 6.86 (5.58) and 7.22 (5.42) years for males and females, respectively. **Figure 2** shows the correlation between chronological age (y-axis) and predicted brain age (x-axis)^28^ in the out-of-sample control (males r=0.87, p<0.001; R²=0.76 and females r=0.86, p<0.001; R²=0.74), and MDD test samples (males r=0.81, p<0.001; R^2^=0.66 and females r=0.82, p<0.001; R^2^=0.68). The model also showed relatively good generalization to completely independent healthy control samples of the ENIGMA Bipolar Disorder working group (MAE=7.24 [SD 5.82]; r=0.76, p<0.001; R^2^=0.57 for males and MAE=7.45 [5.44]; r=0.75, p<0.001; R^2^=0.56, for females), **appendix**.

**Figure 2:**
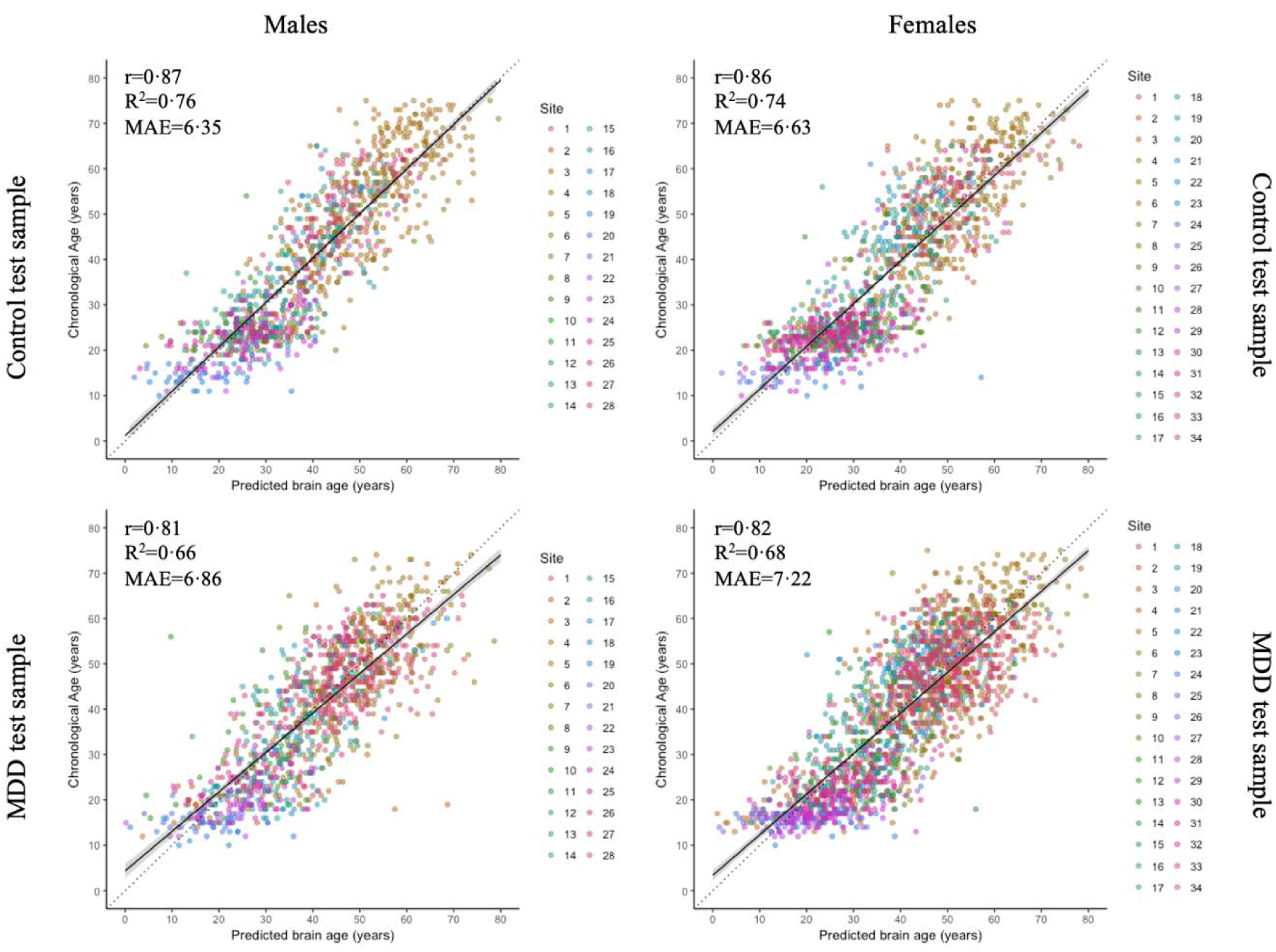
Brain age prediction based on 7 FreeSurfer subcortical volumes, lateral ventricles, 34 cortical thickness and 34 surface area measures, and total intracranial volume. The plots show the correlation between chronological age and predicted brain age in the test samples, derived from the 10-fold cross-validation of the Support Vector Regression model in the training samples, separately for males (*left*) and females (*right*). The colors indicate scanning sites and each circle represents an individual subject: the *upper panels* display controls and the *lower panels* MDD patients. Diagonal dashed line reflects the line of identity (x=y).

### MDD patients show increased brain-PAD compared to controls

There was a main effect of diagnostic group. Specifically, individuals with MDD showed +0.90 (SE 0.21) years higher brain-PAD than controls (p<0.0001, Cohen’s d=0.12, 95% CI 0.06-0.17), **figure 3**. Additionally, we found significant main effects for age, age^2^, and age^3^ (b=-0.02-0.72, all p’s<0.0001), and a trend for a main effect of sex, with higher brain-PAD in females (b=0.39, p=0.0501). Our analyses revealed no significant three-way interaction between diagnosis-by-age-by-sex, nor significant two-way interactions. Of note, there were no significant interactions with age, age^2^, or age^3^ and MDD status; thus, the residual age effects in the brain-PAD estimates did not influence the case-control difference. Further, all nonlinear age effects were accounted for in analyses. All FreeSurfer features, except the entorhinal and temporal pole average thickness, showed a significant (P_FDR_<0.05) association with brain-PAD. All features, except the mean lateral ventricles, yielded negative associations, and are visualized in **figure 4**.

**Figure 3:**
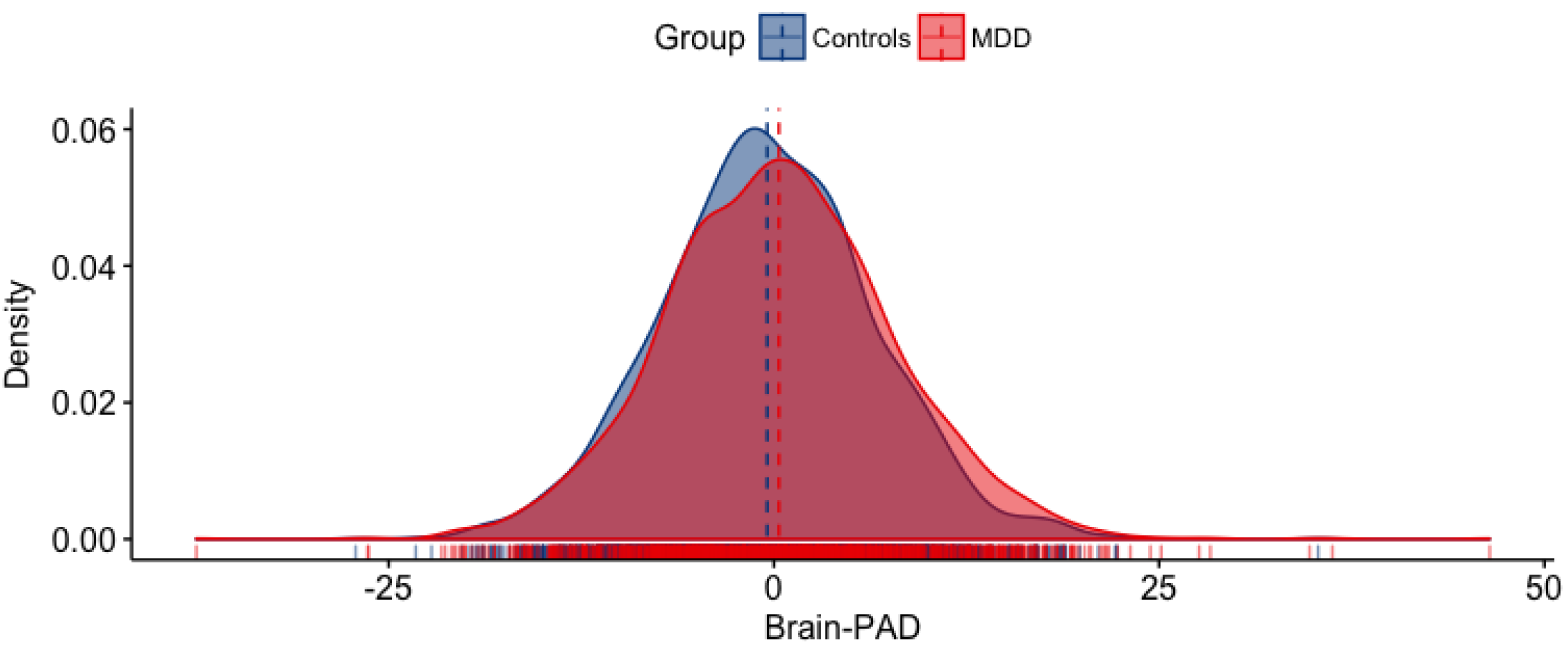
Case-control differences in brain aging. Brain-PAD (predicted brain age - chronological age) in patients with major depressive disorder (MDD) and controls. Group level analyses showed that MDD patients exhibited significantly higher brain-PAD than controls (b=0.90, p<0.0001), although large within-group variation and between-group overlap is observed. The brain-PAD estimates are adjusted for chronological age, age^2^, age^3^, sex and scanning site.

**Figure 4:**
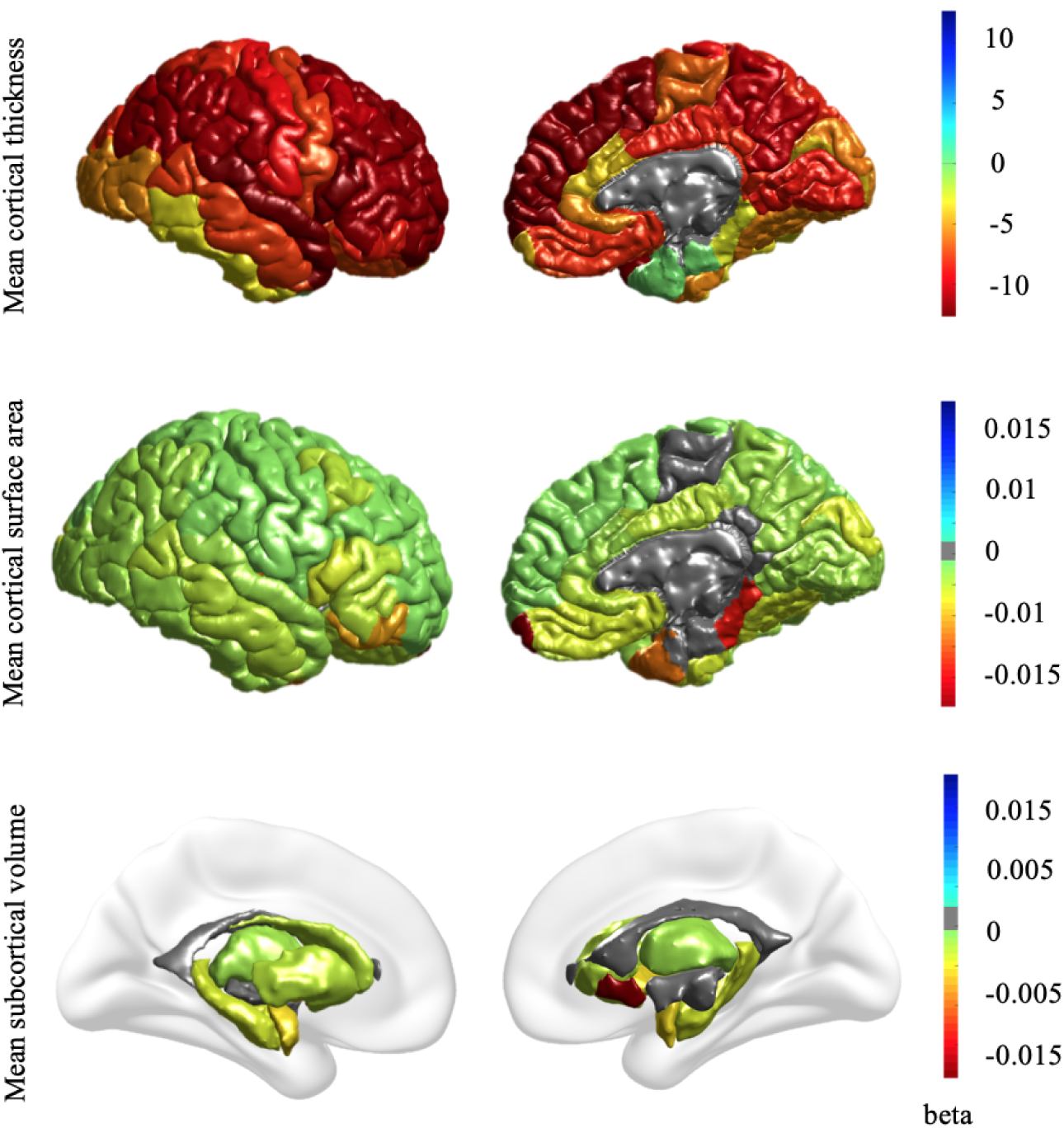
Univariate associations between brain predicted age difference (predicted brain age - chronological age; brain-PAD) and FreeSurfer measures across controls and major depressive disorder (MDD) groups. Effect sizes (regression coefficients) are shown for regions with a significant (P_FDR_<0.05) negative association with brain-PAD, only the mean lateral ventricles yielded a significant positive association. The figure shows associations with cortical thickness measures (*top row*), cortical surface areas (*middle row*), and subcortical volumes (*bottom row*). The brain-PAD estimates are adjusted for chronological age, age^2^, age^3^, sex and scanning site. The significant negative association with ICV was excluded from this figure for display purposes.

### Clinical characteristics and brain-PAD

Strongest effects of higher brain-PAD were observed in patients with late age of onset of depression (>55 years; +1.7 years, *p*=0.009, Cohen’s *d*=0.17), currently depressed (+1.2y, *p*<0.0001, *d*=0.13), and first episode (+1.2y, *p*=0.0001, *d*=0.12) MDD patients, compared to controls. However, we observed relatively similar effects in remitted (+1.2y, p=0.01, d=0.11), both antidepressant users and antidepressant medication-free (both +0.9y, p’s<0.002, d=0.09), early age of onset of depression (<26 years; +0.8y, p=0.0005, d=0.10), and recurrent depressed patients (+0.7y, p=0.003, d=0.08), as well as in those with an adult age of onset of MDD (+0.5y, p=0.02, d=0.06), compared to controls (**table 1**). Post-hoc comparisons between the MDD subgroups did not show any significant differences (i.e., first vs. recurrent episode, antidepressant medication-free vs. antidepressant users, remitted vs. currently depressed patients, or early vs. adult vs. late age of onset of depression). Brain-PAD was positive in all MDD subgroups, and there were no negative associations with any clinical characteristics.

**Table 1:**
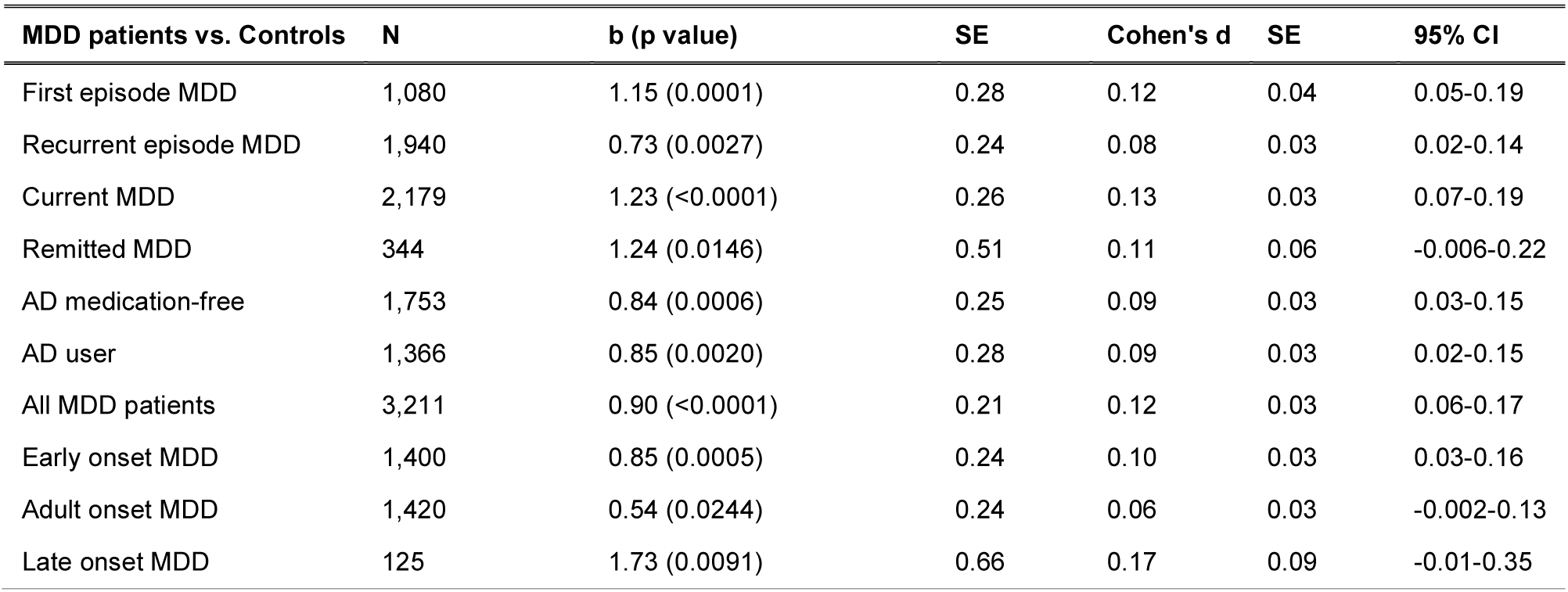
Clinical characteristics and brain aging. Positive brain-PAD scores (predicted brain age - chronological age) were found for all subgroups of patients with major depressive disorder (MDD) compared to controls (N=2,256). b=regression coefficient; this indicates the average brain-PAD difference between MDD patients and controls in years. AD, Antidepressant.

### Increased brain-PAD is associated with greater depressive symptom severity

There was an association with depression severity at the time of scanning within the MDD sample, illustrated by higher brain-PAD in individuals with more severe self-reported depressive symptomatology (b=0.05, p=0.004) as measured in N=1,538 patients who completed the BDI-II. We were not able to confirm this, however, in N=1,905 depressed individuals who were assessed using the HDRS-17 clinician-based questionnaire (b=0.003, p=0.90).

## Discussion

Using a brain age algorithm based on commonly used brain measures derived from T1-weighted scans from over 3,500 males and 4,900 females, we found subtle age-associated gray matter differences in major depressive disorder (MDD). At the group level, the brain age model predicted chronological age in controls and MDD patients from 77 brain morphometric features, and patients had, on average, a 0.90 years greater discrepancy between their predicted and actual age compared to control participants. Strongest effects were observed in late-life onset of depression (+1.7y, d=0.17), currently depressed (+1.2y, d=0.13), and first episode MDD (+1.2y, d=0.12) patients, compared to controls. Finally, each one-point increase in self-reported symptom severity score at study inclusion added, on average, 18 days of brain aging, potentially underscoring the importance of reducing the number of symptoms in the treatment of depression.

The positive association between brain aging and symptom severity, measured with the self-report BDI-II questionnaire, was not confirmed using the clinician-based HDRS-17. Post-hoc analyses in overlapping samples with both scores (N=1,302) yielded a significant correlation between them (r=0.67, p<0.0001), yet the same discrepant association with brain-PAD. This could perhaps be explained by the differential proportion of items emphasizing cognitive and affective (BDI-II) or somatic and behavioral dimensions (HDRS-17).^29^ Alternatively, brain age may be more sensitive to subjective (BDI) than to objectively (HDRS-17) rated experiences, consistent with the finding of Kwak and colleagues (2018) that the subjective experience of aging was closely related to predicted brain age.^30^ However, it is important to bear in mind the small effect size (b=0.05). Nonetheless, positive associations with current depressive symptom severity have been previously reported with more advanced levels of biological aging, as indicated by shorter telomere length^31^ and increased epigenetic aging.^19^

This study showed relatively largest effect size of advanced brain aging in patients with a late-life onset of depression (>55 years old) compared to controls. However, we did not find significant differences between early vs. adult vs. late onset of depression groups. Additionally, no differences between remitted (N=344) and acute patients (N=2,179) were found, leading to the speculation that an initial brain insult during a first episode of depression or preceding clinical disease onset may leave a lasting impact even after remission. To date, the reversibility of gray matter alterations in MDD over time remains rather elusive due to the lack of reliable longitudinal studies.^32^ Yet, cross-sectional studies show that “younger” appearing brains are seen in groups of individuals with greater physical activity,^33^ long-term meditation practitioners,^11^ and amateur musicians,^34^ suggesting that brain age might be a modifiable metric. Moreover, one study suggests dynamic potential by showing that in healthy individuals brain-PAD was temporarily reduced by 1.1 years due to the probable acute anti-inflammatory effects of ibuprofen.^35^ In this study, there was no detectable effect of antidepressant use on brain aging within MDD individuals. As antidepressants are suggested to exert a neuroprotective effect, for example by promoting brain-derived neurotrophic factor (BDNF),^36^ it remains to be elucidated how adaptable brain age is in response to pharmacotherapy. However, the cross-sectional nature of the current study and the lack of detailed information on lifetime use, dosage and duration of use of antidepressants, do not allow us to draw any conclusions regarding direct effects of antidepressants on brain aging. Thus, longitudinal research and randomized controlled intervention studies are needed to develop an understanding of how reversible brain aging is after remission of MDD and how modifiable in response to pharmacology, but also to non-pharmacological strategies (e.g., psychological, exercise and/or nutritional interventions), as seen in other biological age indicators.^37–39^

Further, the currently observed effect size of Cohen’s d=0.12 with regard to brain aging is consistent with previously seen modest structural brain differences in MDD. Earlier work from the ENIGMA MDD working group also showed small subcortical (hippocampus; d=-0.14), and small to moderate cortical reductions (e.g. left medial orbitofrontal cortex thickness in adults, d=-0.13 and right lingual gyrus surface area in adolescents, d=-0.42) in patients compared to controls.^15,16^ Here, we particularly find strong widespread significant negative associations between brain aging and cortical thickness, and comparably weaker associations with surface area and subcortical volume measures (**figure 4**), consistent with literature on age-related structural brain changes in adolescents^40^ and adults.^41^ We also visualized these associations separately for controls and MDD patients, but findings were similar and suggest comparable spatial brain aging patterns in both groups (**appendix**). Notably, we did not include a spatial weight map of our brain age model, as the weights (although linear) are obtained from a multivariable model, and do not allow for a straightforward interpretation of the importance of the brain regions contributing to the aging pattern.

Our findings were in contrast to earlier work showing a +4.0 years of brain aging in a smaller sample of MDD patients (N=104; 18-65 years).^6^ However, a recent preliminary study in 211 MDD patients (18-71 years) found a similar effect size to ours, albeit non-significant (d=0.10, p=0.33).^26^ In the latter study, brain-PAD was derived using a brain age model trained on >12,000 healthy individuals (vs. the 800 in the Koutsouleris study^6^ vs. >1,100 in this study), emphasizing the relevance of sample size for both training and test samples for sensitivity to detect reliable, yet subtle, effects. Similarly, with respect to reaching statistical significance, large sample sizes are needed to detect small effect sizes commonly found with biological age indicators,^18,19,31^ but also other markers (e.g. BDNF, cortisol, oxidative stress)^42–44^ in depression research. A major strength of this study is, therefore, the mega-analytic approach of pooling harmonized data from many heterogeneous sites, making predictive models less susceptible to overfitting^45^ and more generalizable to other populations.^46^

Inflammation may be a common biological mechanism between MDD and brain aging. Neuroimmune mechanisms (e.g. pro-inflammatory cytokines) influence biological processes (e.g. synaptic plasticity), and inflammatory biomarkers are commonly dysregulated in depression.^47^ Both cerebrospinal fluid and peripheral blood interleukin (IL)-6 levels are elevated in MDD,^48^ and increased IL-6 expression may affect brain morphology through neurodegenerative processes.^49^ Moreover, work by Kakeda and colleagues (2018) demonstrated a significant inverse relationship between IL-6 levels and surface-based cortical thickness and hippocampal subfields in medication-free, first-episode MDD patients.^50^ This accords with the current observation of increased brain-PAD in medication-free and first-episode patients, compared to controls, perhaps suggesting that neuroimmune mechanisms may be chief candidates involved in the brain morphology alterations, also in the early stage of illness. Further, the age-related structural alterations in MDD may also be explained by shared underlying (epi)genetic mechanisms involved in brain development and plasticity (thereby influencing brain structure) and psychiatric illness.^51^ For instance, Aberg and colleagues (2018) showed that a significant portion of the genes represented in overlapping blood-brain methylome-wide association findings for MDD were important for brain development, such as induction of synaptic plasticity by BDNF.^52^

Our current findings in MDD show lower brain aging than previously observed in schizophrenia (SCZ) (brain-PAD ranges from +2.6 – +5.5y, d=0.64)^6,22^, even in early stages of first episode SCZ.^25^ Inconsistent findings are reported in bipolar disorder (BD), with “younger” brain age^23^ or no differences compared to controls.^25^ However, more studies with larger sample sizes are needed to confirm brain aging in these psychiatric disorders-endeavors currently pursued by other ENIGMA psychiatric disease working groups using the same brain age models, which will allow future cross-disorder comparisons between brain-PAD in e.g. MDD, BD and SCZ.

While our results are generally consistent with existing literature on advanced or premature biological aging and major depression using other biological indicators,^18^ it is important to critically consider the current findings and note their limitations. First, limited information was available on clinical characterization and brain-PAD could not be compared against somatic health outcomes here. Second, given the relatively crude and limited number of gray matter features, the best MAE that could be achieved was 6.9 years, compared to ∼4.9 years accomplished by other brain age predictors (e.g., those based on spatial images with high dimensional features that may also include white matter).^12^ However, an advantage to using FreeSurfer data over voxelwise methods is that the fewer dimensions render our models less prone to overfitting and more flexible in exploring the use of different machines and kernels (**appendix**). Furthermore, pooling data from many scanning sites comes at the cost of increasing heterogeneity of MRI data and other sample specifics. However, withstanding the latter limitation, models are therefore consequently tested on “ecologically valid” samples, bolstering confidence in their deployability and shareability.^53^ Finally, the large within-group variance regarding the brain-PAD outcome in both controls and MDD (**figure 3**), compared to the small between-group variance, renders the use of this brain aging indicator for discriminating patients and controls at the individual level difficult. As many of the MDD patients do not show advanced brain aging compared to controls, the clinical significance of the observed higher brain-PAD in MDD patients in this study may be limited. Yet, interindividual differences highlight the importance of studying the individual, rather than the average patient^54^ and provide the opportunity to elucidate whether a subgroup of patients with high brain-PAD may be at risk for worse psychiatric, neurologic, and somatic health outcomes. Local sites that participated in this study with clinical phenotyping and longitudinal information on mental and somatic health outcomes (e.g., genomic variation, omics profiles, comorbidities, lifestyle, inflammation, oxidative stress, chronic diseases) will allow further evaluation of the predictive value of the brain-PAD estimates. This is expected to promote continued growth of knowledge in pursuance of useful clinical applications.

In conclusion, compared to controls, both male and female MDD patients show advanced brain aging, with a subtle association with current symptom severity. This is consistent with other studies of biological aging indicators in MDD at cellular and molecular levels of analysis (i.e., telomere length and epigenetic age). The deviation of brain metrics from normative aging trajectories in MDD may contribute to increased risk for mortality and aging-related diseases commonly seen in MDD. However, the substantial within-group variance and overlap between groups signify that more (longitudinal) work including in-depth clinical characterization and more precise biological age predictor systems are needed to elucidate whether brain age indicators can be clinically useful in MDD. Future studies may use our current ENIGMA brain age prediction model to associate brain-PAD with treatment response and other available information on longitudinal mental and somatic health outcomes, other aging indicators, and incidence and/or prevalence of other chronic diseases in their local samples in pursuance of greater clinical application.

## Authors contributions

### Concept and design

AFM, BP, JHC, LKMH, LS, LTE, NJ, PMT.

### Acquisition, analysis or interpretation of data

AA, AC, AFM, AHS, AJ, AK, AMM, ANS, AS, AU, BAM, BCD, BG, BH, BJH, BJO, BK, BKD, BL, BP, BTB, CA, CC, CF, CGC, CGD, CH, CK, CM, CMB, CMD, DD, DG, DHW, DJS, DMC, EB, ECR, EF, EO,EPC, ES, EV, FLSD, FMH, FPM, GBF, GBH, GdZ, GM, GR, GT, GZ, HCW, HGR, HJG, HST, HV,HW, IB, IBH,IHG, IMV, JH, JHC,JL, JMF, JMG, JR, JR, JS, JS, JS, KB, KC, KLM, KS, KS, KW, LA, LKMH, LN, LR, LS, LTE, LTS, MA, MA, MA, MCGO,MDS, MGSS, MH, MHS, MI, MJP, MJvT, MJW, ML, MMR, MP, MVZ, NG, NH, NRW, NW, OAA, OG, OS, PBM, PF, PGPR, PGS, PMT, PRS, RD, RK, RL, RM, RS, RV, SF, SF, SIT, SNH, SSM, SW, TCH, TDS, TE, TF, TH, TH, TK, TMC, TTY, UD, UFM, VE, VZ, XC.

### Drafting of the manuscript

LKMH, LS.

### Critical revision of the manuscript for important intellectual content

AA, AC, AFM, AHS, AMM, AS, AU, BAM, BCD, BH, BJH, BK, BKD, BM, BP, BTB, CA, CC, CF, CGC, CGD, CK, CMB, DG, DJS, EB, EP, ES, EV, FLSD, FPM, GBF, GBH, GZ, HCW, HGR, HJG, HST, HV, HW, IHG, IMV, JHC, JMG, JR, JR, JS, JS, KB,KC, KLM, KS, KW, LA, LKMH, LR, LS, LTE, LTS, MA, MA, MA, MDS, MH, MJP, MJW, ML, MMR, MW, NG, NH, NJ, NRW, NW, OAA, OG, PGS, PMT, RD, RK, RV, SEM, SF, SF, SW, TCH, TE, TF, TH, TH, TTY, UD, UFM, VE, XC.

### Statistical analysis

AFM, JHC, LKMH, LS, RD.

### Obtained funding

AJ, AK, AMM, BH, BJH, BL, BP, CGD, CH, CMB, CMD, DMC, EB, EPC, EV, FMH, FPM, GBF, GZ, HJG, HV, IBH, IMV, JH, JMF, JS, KB, KLM, KS, KS, LR, LS, MA, MCGO, MGSS, MI, MJW, ML, MP, NG, NH, NJ, OAA, OG, PBM, PMT, PRS, RM, RS, TE, TF, TH, TK, UD, UFM, XC.

### Administrative, technical or material support

AA, AHS, AS, AU, BCD, BH, BL, BM, BP, CA, CMB, CMD, DJS, DMC, ECR, EP, EPC, EV, FMH, GR, GZ, HGR, HJG, HST, HV, IMV, JMF, JMG, JR, JS, KLM, KS, LN, LR, LTS, MA, MCGO, MGSS, MJW, MMR, MW, NG, NH, NJ, OAA, PBM, PGS, PMT, PRS, RM, RS, SEM, SF, SSM, TH, XC.

Supervision: AFM, BP, JHC, LS.

All authors approved the content of the manuscript.

## Acknowledgments

ENIGMA MDD: This work was supported by NIH grants U54 EB020403 and R01 MH116147.

BiDirect-Münster: The study was supported by a grant from the German Federal Ministry of Education and Research (BMBF; grant FKZ-01ER0816 and FKZ-01ER1506).

Calgary: This study was supported by the Alberta Children's Hospital Foundation.

CliNG (Heidelberg): This work was partially supported by the Deutsche Forschungsgemeinschaft (DFG) via grants to OG (GR1950/5-1 and GR1950/10-1).

CODE: The CODE cohort was collected from studies funded by Lundbeck and the German Research Foundation (WA 1539/4-1, SCHN 1205/3-1, SCHR443/11-1).

DIP-Groningen: This study was supported by the Gratama Foundation, the Netherlands (2012/35 to NG)

Dublin: The study was funded by Science Foundation Ireland, with a Stokes Professorship Grant to TF.

Edinburgh: The research leading to these results was supported by IMAGEMEND, which received funding from the European Community's Seventh Framework Programme (FP7/2007-2013) under grant agreement no. 602450. This paper reflects only the author’s views and the European Union is not liable for any use that may be made of the information contained therein. This work was also supported by a Wellcome Trust Strategic Award 104036/Z/14/Z.

FOR2107-Marburg: This work was funded by the German Research Foundation (DFG, grant FOR2107 KR 3822/7-2 to AK; FOR2107 KI 588/14-2 to TK and FOR2107 JA 1890/7-2 to AJ).

Leiden: EPISCA was supported by GGZ Rivierduinen and the LUMC.

Melbourne: This study was funded by National Health and Medical Research Council of Australia (NHMRC) Project Grants 1064643 (Principal Investigator BJH) and 1024570 (Principal Investigator CGD).

Minnesota: This study was funded by the National Institute of Mental health grant K23MH090421 (D. Cullen) and Biotechnology Research Center grant P41RR008079 (Center for Magnetic Resonance Research), the National Alliance for Research on Schizophrenia and Depression, the University of Minnesota Graduate School, and the Minnesota Medical Foundation. This work was carried out in part using computing resources at the University of Minnesota Supercomputing Institute.

Münster: This work was funded by the German Research Foundation (DFG, grant FOR2107 DA1151/5-1 and DA1151/5-2 to UD; SFB-TRR58, Projects C09 and Z02 to UD) and the Interdisciplinary Center for Clinical Research (IZKF) of the medical faculty of Münster (grant Dan3/012/17 to UD).

Novosibirsk: This work was supported by Russian Science Foundation (RSF grant 16-15-00128) to LA.

SHIP: The Study of Health in Pomerania (SHIP) is part of the Community Medicine Research net (CMR) (http://www.medizin.uni-greifswald.de/icm) of the University Medicine Greifswald, which is supported by the German Federal State of Mecklenburg-West Pomerania. MRI scans in SHIP and SHIP-TREND have been supported by a joint grant from Siemens Healthineers, Erlangen, Germany and the Federal State of Mecklenburg-West Pomerania. This study was further supported by the EU-JPND Funding for BRIDGET (FKZ:01ED1615).

Stanford: This work was supported by NIH grant R37 MH101495.

Sydney: This study was supported by the following National Health and Medical Research Council funding sources: Programme Grant (no. 566529), Centres of Clinical Research Excellence Grant (no. 264611), Australia Fellowship (no. 511921) and Clinical Research Fellowship (no. 402864).

The QTIM dataset was supported by the Australian National Health and Medical Research Council (Project Grants No. 496682 and 1009064) and US National Institute of Child Health and Human Development (RO1HD050735).

Geraldo Busatto was supported by the funding agencies FAPESP and CNPq, Brazil.

Christopher Ching was supported by NIH grants U54 EB020403, RF1 AG041915, RF1AG051710, P41EB015922, R01MH116147, and R56AG058854.

James Cole was funded by a UKRI Innovation Fellowship.

Baptiste Couvy-Duchesne was supported by a NHMRC CJ Martin Fellowship (APP1161356).

Cynthia Fu was supported in part by MRC grant, NIHR BRC grant.

Beata Godlewska was supported by the Medical Research Council.

Tiffany Ho was supported by the National Institute of Health (K01MH117442).

Neda Jahanshad was supported by NIH grants R01 MH117601, R01 AG059874, U54 EB020403, RF1 AG041915, RF1AG051710, P41EB015922, R01MH116147, and R56AG058854.

Andre Marquand was supported by the Dutch Organization of Scientific Research under a Vernieuwingsimpuls 'VIDI' Fellowship (grant number 016.156.415)

Sarah Medland was supported by an Australian National Health and Medical Research Council Senior Research Fellowship (APP1103623).

Maria Portella was funded by Ministerio de Ciencia e Innovación of Spanish Government (ISCIII) through a "Miguel Servet II" (CP16/00020).

Philipp Sämann reports funding by the German Research Foundation (DFG, SA 1358/2-1) and the Max Planck Institute of Psychiatry, Munich.

Lianne Schmaal was supported by a NHMRC Career Development Fellowship (1140764). Jair Soares was supported by the Pat Rutherford Chair in Psychiatry, UTHealth.

Paul Thompson was supported in part by NIH grants U54 EB020403, RF1 AG041915, RF1AG051710, P41EB015922, R01MH116147, and R56AG058854.

Sophia Thomopoulos was supported in part by NIH grants U54 EB020403, RF1 AG041915, RF1AG051710, P41EB015922, R01MH116147, and R56AG058854.

Tony Yang was supported for this study by: NIMH R01MH085734, NCCIH R21AT009173, UCSF Research Evaluation and Allocation Committee (REAC) and J. Jacobson Fund, the Brain and Behavior Research Foundation (formerly NARSAD).

Amsterdam DIADE: The DIADE study was funded by ZonMW OOG 2007, the Netherlands (#100002034).

Cardiff: The Cardiff dataset was supported through a 2010 NARSAD Young Investigator Award (ref: 17319) to XC.

CIAM Cape Town: This work was supported by the University Research Council of the University of Cape Town and the National Research Foundation of South Africa.

FIDMAG Barcelona: This work was supported by the Generalitat de Catalunya (2014 SGR 1573) and Instituto de Salud Carlos III (CPII16/00018) and (PI14/01151 and PI14/01148).

Galway: This work was supported by the Health Research Board, Ireland and the Irish Research Council. Grenoble: This work was supported by research grants from Grenoble University Hospital.

Halifax: This work was supported by the Canadian Institutes of Health Research (142255).

MOODINFLAME Groningen: This study was funded by EU-FP7-HEALTH-222963 ‘MOODINFLAME’ and EU-FP7-PEOPLE-286334 ‘PSYCHAID’.

Oslo: Funded by the South-Eastern Norway Regional Health Authority (2014097) and a research grant from Mrs. Throne-Holst.

Paris: This work was supported by the FRM (Fondation pour la recherche Biomédicale) "Bio-informatique pour la biologie" 2014 grant.

Singapore: Funded by Singapore Bioimaging Consortium Research Grant (SBIC RP C-009/2006) and NHG grant (SIG/15012).

UNSW: Australian NHMRC Program Grant 1037196 and Project Grants 1063960 and 1066177; and the Janette Mary O’Neil Research Fellowship to JMF.

VA San Diego Healthcare/University of California San Diego: This study was supported by R01MH083968, Desert-Pacific Mental Illness Research Education and Clinical Center, and the US National Science Foundation (Science Gateways Community Institutes; XSEDE).

Ole Andreassen was funded by the Research Council of Norway (223273, 248778, 273291), NIH (ENIGMA grants).

Caterina Bonnin thanks the PERIS grant contract by Departament de Salut CERCA Programme/Generalitat de Catalunya SLT002/16/00331.

Jose Goikolea thanks the support of CIBERSAM and the Comissionat per a Universitats i Recerca del DIUE de la Generalitat de Catalunya to the Bipolar Disorders Group (2017 SGR 1365) and the project SLT006/17/00357, from PERIS 2016-2020 (Departament de Salut). CERCA Programme/Generalitat de Catalunya.

Tomas Hajek was supported by the Canadian Institutes of Health Research (103703, 106469), Nova Scotia Health Research Foundation, Dalhousie Clinical Research Scholarship, Brain & Behavior Research Foundation (formerly NARSAD) 2007 Young Investigator and 2015 Independent Investigator Awards.

Mikael Landén was funded by the Swedish state under the ALF-agreement (ALF 20170019, ALFGBG-716801) and the Swedish Research Council (2018-02653).

Joaquim Radua thanks the Miguel Servet contract by the Spanish Ministerio de Ciencia, Innovacion y Universidades.

Jonathan Savitz was supported by the National Institute of General Medical Sciences (P20GM121312) and the National Institute of Mental Health (R21MH113871)

Mauricio Seroa was supported by the funding agencies CAPES, Brazil.

Dan Stein was supported by the SAMRC.

Garrett Timmons’ work was supported by the National Institutes of Health, Grant T35 AG026757/AG/NIA and the University of California San Diego, Stein Institute for Research on Aging.

Eduard Vieta thanks the support of the Spanish Ministry of Science, Innovation and Universities (PI15/00283) integrated into the Plan Nacional de I+D+I y cofinanciado por el ISCIII-Subdirección General de Evaluación y el Fondo Europeo de Desarrollo Regional (FEDER); CIBERSAM; and the Comissionat per a Universitats i Recerca del DIUE de la Generalitat de Catalunya to the Bipolar Disorders Group (2017 SGR 1365) and the project SLT006/17/00357, from PERIS 2016-2020 (Departament de Salut). CERCA Programme/Generalitat de Catalunya.

Marcus Zanetti was supported by FAPESP, Brazil (grant no. 2013/03905-4).

## Conflicts of interest

These authors all declare no conflicts of interest:

Lyubomir Aftanas, Moji Aghajani, André Aleman, Bernhard Baune, Klaus Berger, Ivan Brak, Geraldo Busatto Filho, Angela Carballedo, Christopher Ching, James Cole, Colm Connolly, Baptiste Couvy-Duchesne, Kathryn Cullen, Udo Dannlowski, Christopher Davey, Danai Dima, Richard Dinga, Fabio Duran, Verena Enneking, Lisa Eyler, Elena Filimonova, Stefan Frenzel, Thomas Frodl, Cynthia Fu, Beata Godlewska, Ian Gotlib, Nynke Groenewold, Dominik Grotegerd, Oliver Gruber, Tim Hahn, Geoffrey Hall, Laura Han, Ben Harrison, Sean Hatton, Marco Hermesdorf, Tiffany Ho, Norbert Hosten, Neda Jahanshad, Andreas Jansen, Claas Kähler, Tilo Kircher, Bonnie Klimes-Dougan, Bernd Krämer, Axel Krug, Jim Lagopoulos, Ramona Leenings, Frank MacMaster, Glenda MacQueen, Andre Marquand, Andrew McIntosh, Katie McMahon, Sarah Medland, Philip Mitchell, Bryon Mueller, Benson Mwangi, Evgeny Osipov, Maria Portella, Elena Pozzi, Liesbeth Reneman, Jonathan Repple, Pedro Rosa, Matthew Sacchet, Philipp Sämann, Lianne Schmaal, Anouk Schrantee, Egle Simulionyte, Jens Sommer, Dan Stein, Olaf Steinsträter, Lachlan Strike, Sophia Thomopoulos, Marie-José van Tol, Ilya Veer, Robert Vermeiren, Henrik Walter, Nic van der Wee, Steven van der Werff, Heather Whalley, Nils Winter, Katharina Wittfeld, Margaret Wright, Mon-Ju Wu, Dick Veltman, Henry Völzke, Tony Yang, Vasileios Zannias, Greic de Zubicaray, Giovana Zunta-Soares, Christoph Abé, Martin Alda, Ole Andreassen, Erlend Bøen, Caterina Bonnin, Erick Canales-Rodriguez, Dara Cannon, Xavier Caseras, Tiffany Chaim-Avancini, Pauline Favre, Sonya Foley, Janice Fullerton, Jose Goikolea, Bartholomeus Haarman, Tomas Hajek, Chantal Henry, Josselin Houenou, Fleur Howells, Martin Ingvar, Rayus Kuplicki, Beny Lafer, Rodrigo Macha-Vieira, Ulrik Malt, Colm McDonald, Philip Mitchell, Leila Nabulsi, Maria Concepcion Garcia Otaduy, Bronwyn Overs, Mircea Polosan, Edith Pomarol-Clotet, Joaquim Radua, Maria Rive, Gloria Roberts, Henricus Ruhe, Raymond Salvador, Salvador Sarró, Theodore Satterthwaite, Jonathan Savitz, Aart Schene, Peter Schofield, Mauricio Serpa, Kang Sim, Marcio Gerhardt Soeiro-de-Souza, Ashley Sutherland, Henk Temmingh, Garrett Timmons, Anne Uhlmann, Daniel Wolf, Marcus Zanetti.

These authors received the following funding, however, all unrelated to the current manuscript: Beata Godlewska has received a (non-related) travel grant from Janssen UK. Hans Grabe has received travel grants and speakers’ honoraria from Fresenius Medical Care and Janssen Cilag. He has received research funding from the German Research Foundation (DFG), the German Ministry of Education and Research (BMBF), the DAMP Foundation, Fresenius Medical Care, the EU "Joint Programme Neurodegenerative Disorders (JPND) and the European Social Fund (ESF)".

Ian Hickie was an inaugural Commissioner on Australia’s National Mental Health Commission (2012-2018). He is the Co-Director, Health and Policy at the Brain and Mind Centre (BMC) University of Sydney. The BMC operates an early-intervention youth services at Camperdown under contract to headspace. Professor Hickie has previously led community-based and pharmaceutical industry-supported (Wyeth, Eli Lily, Servier, Pfizer, AstraZeneca) projects focused on the identification and better management of anxiety and depression. He was a member of the Medical Advisory Panel for Medibank Private until October 2017, a Board Member of Psychosis Australia Trust and a member of Veterans Mental Health Clinical Reference group. He is the Chief Scientific Advisor to, and an equity shareholder in, Innowell. Innowell has been formed by the University of Sydney and PwC to deliver the $30m Australian Government-funded ‘Project Synergy’. Project Synergy is a three-year program for the transformation of mental health services through the use of innovative technologies.

Brenda Penninx has received (non-related) research funding from Boehringer Ingelheim and Jansen Research

Knut Schnell has consulted for Roche Pharmaceuticals and Servier Pharmaceuticals

Jair Soares has received research support from BMS, Forest, Merck, Elan, Johnson & Johnson and COMPASS in the form of grants and clinical trials. He is a member of the speakers’ bureaus for Pfizer, Abbott and Sonify and he is a consultant for Astellas.

Torbjørn Elvsåshagen has served as a speaker for Lundbeck.

Mikael Landén declares that, over the past 36 months, he has received lecture honoraria from Lundbeck pharmaceutical. No other equity ownership, profit-sharing agreements, royalties, or patent.

Paul Thompson has received (non-related) research funding from Biogen, Inc. (Boston).

Eduard Vieta has received grants and served as consultant, advisor or CME speaker for the following entities: AB-Biotics, Abbott, Allergan, Angelini, AstraZeneca, Bristol-Myers Squibb, Dainippon Sumitomo Pharma, Farmindustria, Ferrer, Forest Research Institute, Gedeon Richter, Glaxo-Smith-Kline, Janssen, Lundbeck, Otsuka, Pfizer, Roche, SAGE, Sanofi-Aventis, Servier, Shire, Sunovion, and Takeda.

## References

1 John A, Patel U, Rusted J, Richards M, Gaysina D. Affective problems and decline in cognitive state in older adults: a systematic review and meta-analysis. Psychol Med 2018; : 1–13.

2 Penninx BWJH. Depression and cardiovascular disease: Epidemiological evidence on their linking mechanisms. Neurosci Biobehav Rev 2016. DOI:10.1016/j.neubiorev.2016.07.003.

3 Walker ER, McGee RE, Druss BG. Mortality in mental disorders and global disease burden implications: a systematic review and meta-analysis. JAMA Psychiatry 2015; 72: 334–41.

4 Katon WJ. Epidemiology and treatment of depression in patients with chronic medical illness. Dialogues Clin Neurosci 2011; 13: 7–23.

5 Pfefferbaum A, Rohlfing T, Rosenbloom MJ, Chu W, Colrain IM, Sullivan EV. Variation in longitudinal trajectories of regional brain volumes of healthy men and women (ages 10 to 85 years) measured with atlas-based parcellation of MRI. Neuroimage 2013; 65: 176–93.

6 Koutsouleris N, Davatzikos C, Borgwardt S, et al. Accelerated brain aging in schizophrenia and beyond: A neuroanatomical marker of psychiatric disorders. Schizophr Bull 2014; 40: 1140–53.

7 Kessler RC, Bromet EJ, de Jonge P, Shahly V, Wilcox M. The Burden of Depressive Illness. Public Health Perspectives on Depressive Disorders (2017) 2017; 40. https://books.google.nl/books?hl=en&lr=&id=MOEsDwAAQBAJ&oi=fnd&pg=PT56&dq=burden+major+depression&ots=ZuoTrz61Ow&sig=Lw5ghJk78h50BInYqJcDlLsWnkA.

8 Diniz BS, Vieira EM. Stress, Inflammation, and Aging: An Association Beyond Chance. Am J Geriatr Psychiatry 2018; 26: 964–5.

9 Jylhava J, Pedersen NL, Hagg S. Biological Age Predictors. EBioMedicine 2017; 21: 29–36.

10 Cole JH, Franke K. Predicting Age Using Neuroimaging: Innovative Brain Ageing Biomarkers. Trends Neurosci 2017; 40: 681–90.

11 Luders E, Cherbuin N, Gaser C. Estimating brain age using high-resolution pattern recognition: Younger brains in long-term meditation practitioners. Neuroimage 2016; 134: 508–13.

12 Cole JH, Ritchie SJ, Bastin ME, et al. Brain age predicts mortality. Mol Psychiatry 2017; : 1–8.

13 Liem F, Varoquaux G, Kynast J, et al. Predicting brain-age from multimodal imaging data captures cognitive impairment. Neuroimage 2017; 148: 179–88.

14 Hatton SN, Franz CE, Elman JA, et al. Negative fateful life events in midlife and advanced predicted brain aging. Neurobiol Aging 2018; 67: 1–9.

15 Schmaal L, Veltman DJ, van Erp TGM, et al. Subcortical brain alterations in major depressive disorder: findings from the ENIGMA Major Depressive Disorder working group. Mol Psychiatry 2015; : 1–7.

16 Schmaal L, Hibar DP, Sämann PG, et al. Cortical abnormalities in adults and adolescents with major depression based on brain scans from 20 cohorts worldwide in the ENIGMA Major Depressive Disorder Working Group. Mol Psychiatry 2016; 22: 900.

17 Jahanshad N, Thompson PM. Mini-Review Multimodal Neuroimaging of Male and Female Brain Structure in Health and Disease Across the Life Span. 2017; 379: 371–9.

18 Darrow SM, Verhoeven JE, Révész D, et al. The Association Between Psychiatric Disorders and Telomere Length: A Meta-Analysis Involving 14,827 Persons. Psychosom Med 2016; 78: 776–87.

19 Han LKM, Aghajani M, Clark SL, et al. Epigenetic Aging in Major Depressive Disorder. Am J Psychiatry 2018; : appi.ajp.2018.1.

20 Whalley HC, Gibson J, Marioni R, et al. Accelerated epigenetic ageing in depression. 2017.

21 Cole JH, Marioni RE, Harris SE, Deary IJ, Cole JH. Brain age and other bodily ‘ages’ : implications for neuropsychiatry. Mol Psychiatry 2018. DOI:10.1038/s41380-018-0098-1.

22 Schnack HG, Van Haren NEM, Nieuwenhuis M, Pol HEH, Cahn W, Kahn RS. Accelerated brain aging in schizophrenia: A longitudinal pattern recognition study. Am J Psychiatry 2016; 173: 607–16.

23 Nenadic I, Dietzek M, Langbein K, Sauer H, Gaser C. BrainAGE score indicates accelerated brain aging in schizophrenia, but not bipolar disorder. Psychiatry Research: Neuroimaging 2017. DOI:10.1016/j.pscychresns.2017.05.006.

24 Kolenic M, Franke K, Hlinka J, et al. Obesity, dyslipidemia and brain age in first-episode psychosis. J Psychiatr Res 2018; 99: 151–8.

25 Hajek T, Franke K, Kolenic M, et al. Brain Age in Early Stages of Bipolar Disorders or Schizophrenia. Schizophr Bull 2017; published online Dec 20. DOI:10.1093/schbul/sbx172.

26 Kaufmann T, Meer DVD, Doan NT, et al. Genetics of brain age suggest an overlap with common brain disorders. 2018.

27 Pedregosa F, Varoquaux G, Gramfort A, et al. Scikit-learn: Machine Learning in Python. J Mach Learn Res 2011; 12: 2825–30.

28 Piñeiro G, Perelman S, Guerschman JP, Paruelo JM. How to evaluate models: Observed vs. predicted or predicted vs. observed?. Ecol Modell 2008; 216: 316–22.

29 Brown C, Schulberg HC, Madonia MJ. Assessment depression in primary care practice with the Beck Depression Inventory and the Hamilton Rating Scale for Depression. Psychol Assess 1995. http://psycnet.apa.org/fulltext/1995-27650-001.html.

30 Kwak S, Kim H, Chey J, Youm Y. Feeling How OldI Am: Subjective Age Is Associated With Estimated Brain Age. Front Aging Neurosci 2018; 10: 168.

31 Verhoeven JE, Révész D, Epel ES, Lin J, Wolkowitz OM, Penninx BWJH. Major depressive disorder and accelerated cellular aging: results from a large psychiatric cohort study. Mol Psychiatry 2013; : 1–7.

32 Dohm K, Redlich R, Zwitserlood P, Dannlowski U. Trajectories of major depression disorders: A systematic review of longitudinal neuroimaging findings. Aust N Z J Psychiatry 2017; 51: 441–54.

33 Steffener J, Habeck C, O’Shea D, Razlighi Q, Bherer L, Stern Y. Differences between chronological and brain age are related to education and self-reported physical activity. Neurobiol Aging 2016; 40: 138–44.

34 Rogenmoser L, Kernbach J, Schlaug G, Gaser C. Keeping brains young with making music. Brain Struct Funct 2018; 223: 297–305.

35 Le TT, Kuplicki R, Yeh HW, et al. Effect of Ibuprofen on BrainAGE: A Randomized, Placebo-Controlled, Dose-Response Exploratory Study. Biological Psychiatry: Cognitive Neuroscience and Neuroimaging 2018; : 1–8.

36 Castrén E, Kojima M. Brain-derived neurotrophic factor in mood disorders and antidepressant treatments. Neurobiol Dis 2017; 97: 119–26.

37 Puterman E, Weiss J, Lin J, et al. Aerobic exercise lengthens telomeres and reduces stress in family caregivers: A randomized controlled trial - Curt Richter Award Paper 2018. Psychoneuroendocrinology 2018; 98: 245–52.

38 Chen L, Dong Y, Bhagatwala J, Raed A, Huang Y, Zhu H. Effects of Vitamin D3 supplementation on epigenetic aging in overweight and obese African Americans with suboptimal vitamin D status: a randomized clinical trial. J Gerontol A Biol Sci Med Sci 2018; published online Sept 25. DOI:10.1093/gerona/gly223.

39 Conklin QA, Crosswell AD, Saron CD, Epel ES. Meditation, Stress Processes, and Telomere Biology. Current Opinion in Psychology 2018; published online Nov 19. DOI:10.1016/j.copsyc.2018.11.009.

40 Tamnes CK, Herting MM, Goddings AL. Development of the cerebral cortex across adolescence: A multisample study of interrelated longitudinal changes in cortical volume, surface area and thickness. Journal of 2017. http://www.jneurosci.org/content/early/2017/02/27/JNEUROSCI.3302-16.2017.abstract.

41 Storsve AB, Fjell AM, Tamnes CK, et al. Differential longitudinal changes in cortical thickness, surface area and volume across the adult life span: regions of accelerating and decelerating change. J Neurosci 2014; 34: 8488–98.

42 Black CN, Bot M, Scheffer PG, Cuijpers P, Penninx BWJH. Is depression associated with increased oxidative stress? A systematic review and meta-analysis. Psychoneuroendocrinology 2015; 51: 164–75.

43 Vreeburg S a., Hoogendijk WJG, van Pelt J, et al. Major Depressive Disorder and Hypothalamic-Pituitary-Adrenal Axis Activity. Arch Gen Psychiatry 2009; 66: 617–26.

44 Molendijk ML, Bus BAA, Spinhoven P, et al. Serum levels of brain-derived neurotrophic factor in major depressive disorder: state–trait issues, clinical features and pharmacological treatment. Mol Psychiatry 2011; 16: 1088–95.

45 Yarkoni T, Westfall J. Choosing Prediction Over Explanation in Psychology: Lessons From Machine Learning. Perspect Psychol Sci 2017; 12: 1100–22.

46 Schnack HG, Kahn RS. Detecting neuroimaging biomarkers for psychiatric disorders: Sample size matters. Front Psychiatry 2016; 7. DOI:10.3389/fpsyt.2016.00050.

47 Wohleb ES, Franklin T, Iwata M, Duman RS. Integrating neuroimmune systems in the neurobiology of depression. Nat Rev Neurosci 2016; 17: 497–511.

48 Wang AK, Miller BJ. Meta-analysis of Cerebrospinal Fluid Cytokine and Tryptophan Catabolite Alterations in Psychiatric Patients: Comparisons Between Schizophrenia, Bipolar Disorder, and Depression. Schizophr Bull 2018; 44: 75–83.

49 Frodl T, Carballedo A, Hughes MM, et al. Reduced expression of glucocorticoid-inducible genes GILZ and SGK-1: high IL-6 levels are associated with reduced hippocampal volumes in major depressive disorder. Transl Psychiatry 2012; 2: e88.

50 Kakeda S, Watanabe K, Katsuki A, et al. Relationship between interleukin (IL)-6 and brain morphology in drug-naïve, first-episode major depressive disorder using surface-based morphometry. Sci Rep 2018; 8: 10054.

51 Elliott LT, Sharp K, Alfaro-Almagro F, et al. Genome-wide association studies of brain imaging phenotypes in UK Biobank. Nature 2018; 562: 210–6.

52 Aberg KA, Dean B, Shabalin AA, et al. Methylome-wide association findings for major depressive disorder overlap in blood and brain and replicate in independent brain samples. Mol Psychiatry 2018; published online Sept 21. DOI:10.1038/s41380-018-0247-6.

53 Woo C-W, Chang LJ, Lindquist MA, Wager TD. Building better biomarkers: brain models in translational neuroimaging. Nat Neurosci 2017; 20: 365–77.

54 Wolfers T, Doan NT, Kaufmann T, et al. Mapping the Heterogeneous Phenotype of Schizophrenia and Bipolar Disorder Using Normative Models. JAMA Psychiatry 2018; published online Oct 10. DOI:10.1001/jamapsychiatry.2018.2467.

