## Supplementary_Tables for "Brain Aging in Major Depressive Disorder: Results from the ENIGMA Major Depressive Disorder working group"

**Brain aging in Major Depressive Disorder - Supplementary Tables**

**Content:**

***Supplementary Table S1:*** **ENIGMA - Major Depressive Disorder Working Group Demographics.** Age (in years), and MDD patients-control breakdown for participating sites, separately for males and females, and training and test samples.

***Supplementary Table S2:* ENIGMA - Major Depressive Disorder Working Group Clinical characteristics of MDD patients.** Percentage of MDD patients using antidepressant medication, percentage of first episode and recurrent episode MDD patients, percentage of acutely depressed and remitted MDD patients, age of onset of MDD and severity of symptoms breakdown for participating sites, separately for male and female samples.

***Supplementary Table S3*: ENIGMA - Major Depressive Disorder Working Group Image acquisition and processing by site.**

***Supplementary Table S4*: ENIGMA - Major Depressive Disorder Working Group Instrument for diagnosing Major Depressive Disorder and exclusion criteria by site.**

***Supplementary Table S5:* Alternative machines and kernels in the brain age prediction framework.** Performance metrics in the training samples of males and females across four different machine learning algorithms/kernels are displayed here.

***Supplementary Table S6:* Mean absolute error (MAE) and brain predicted age difference (brain-PAD) per age group in the overall test samples of controls and major depressive disorder (MDD) patients.**

| **Cohort** | | **Sex** | **Training samples** | | **Test samples** | | **Test samples** | |
| --- | --- | --- | --- | --- | --- | --- | --- | --- |
|  |  |  | **Age Controls (Mean ± SD)** | **Total N Controls**  **(N=2,522)** | **Age Controls (Mean ± SD)** | **Total N Controls**  **(N=2,415)** | **Age MDD**  **(Mean ± SD)** | **Total N MDD**  **(N=3,211)** |
| **1** | **Barcelona** | Males | NA | NA | NA | NA | NA | NA |
|  |  | Females | 45·42 ± 8·62 | 12 | 46·27 ± 8·15 | 11 | 46·77 ± 7·95 | 48 |
| **2** | **BiDirect** | Males | 51·42 ± 8·17 | 109 | 51·32 ± 8·18 | 106 | 47·96 ± 7·41 | 231 |
|  |  | Females | 53·10 ± 7·93 | 109 | 52·82 ± 8·10 | 107 | 49·44 ± 7·08 | 341 |
| **3** | **BRDECC London** | Males | 50·20 ± 8·78 | 15 | 52·50 ± 5·35 | 14 | 45·63 ± 10·01 | 22 |
|  |  | Females | 52·65 ± 7·40 | 17 | 51·47 ± 9·99 | 15 | 48·89 ± 8·26 | 47 |
| **4** | **Edinburgh (Bipolar Family Study)** | Males | 23·00 ± 2·52 | 12 | 23·00 ± 2·39 | 8 | 22·29 ± 3·15 | 7 |
|  |  | Females | 22·63 ± 2·44 | 22 | 22·55 ± 2·44 | 20 | 11·82 ± 2·99 | 11 |
| **5** | **Calgary** | Males | 17·50 ± 4·99 | 8 | 17·38 ± 4·69 | 8 | 18·67 ± 2·28 | 30 |
|  |  | Females | 17·83 ± 3·69 | 12 | 17·11 ± 4·96 | 9 | 18·63 ± 2·27 | 30 |
| **6** | **CLiNG** | Males | 25·24 ± 4·81 | 66 | 25·48 ± 5·36 | 64 | 38·70 ± 9·74 | 23 |
|  |  | Females | 25·23 ± 6·11 | 96 | 24·68 ± 4·20 | 95 | 34·19 ± 12·77 | 26 |
| **7** | **CODE (Berlin)** | Males | 48·14 ± 10·70 | 7 | 50·50 ± 11·39 | 4 | 44·40 ± 3·71 | 5 |
|  |  | Females | 32·54 ± 12·89 | 13 | 34·63 ± 12·89 | 8 | 39·62 ± 12·01 | 39 |
| **8** | **Dublin** | Males | 32·23 ± 10·13 | 39 | 31·17 ± 10·42 | 35 | 36·48 ± 10·11 | 40 |
|  |  | Females | 34·24 ± 11·63 | 38 | 32·27 ± 10·48 | 33 | 39·40 ± 11·54 | 48 |
| **9** | **EPISCA (Leiden)** | Males | NA | NA | NA | NA | NA | NA |
|  |  | Females | 14·93 ± 1·82 | 14 | 14·58 ± 1·51 | 12 | 15·50 ± 1·32 | 16 |
| **10** | **FOR2107 - Marburg** | Males | 32·75 ± 11·15 | 60 | 33·12 ± 11·77 | 60 | 37·34 ± 13·56 | 99 |
|  |  | Females | 33·85 ± 13·27 | 98 | 33·14 ± 13·03 | 97 | 38·02 ± 13·58 | 166 |
| **11** | **FOR2107 - Münster** | Males | 26·89 ± 7·62 | 19 | 27·88 ± 10·89 | 17 | 32·43 ± 11·38 | 21 |
|  |  | Females | 24·59 ± 6·34 | 37 | 26·74 ± 11·29 | 34 | 33·62 ± 13·58 | 26 |
| **12** | **Groningen (DIP)** | Males | NA | NA | NA | NA | NA | NA |
|  |  | Females | 41·25 ± 14·18 | 8 | 41·50 ± 14·65 | 8 | 46·50 ± 14·06 | 14 |
| **13** | **Houston** | Males | 37·24 ± 11·85 | 17 | 40·25 ± 11·21 | 16 | 39·83 ± 13·82 | 23 |
|  |  | Females | 37·36 ± 11·97 | 36 | 38·24 ± 13·57 | 34 | 38·67 ± 13·42 | 55 |
| **14** | **Houston adolescents** | Males | 13·05 ± 2·15 | 22 | 13·40 ± 2·06 | 20 | 13·47 ± 2·26 | 15 |
|  |  | Females | 14·47 ± 1·77 | 15 | 14·07 ± 2·27 | 14 | 14·00 ± 2·16 | 10 |
| **15** | **McMaster University Mood Disorders** | Males | 28·40 ± 9·17 | 10 | 29·00 ± 11·31 | 8 | 34·46 ± 13·64 | 24 |
|  |  | Females | 30·56 ± 11·58 | 16 | 30·93 ± 13·66 | 14 | 35·11 ± 13·43 | 27 |
| **16** | **Melbourne** | Males | 19·79 ± 3·19 | 24 | 19·52 ± 2·84 | 21 | 19·21 ± 2·86 | 34 |
|  |  | Females | 19·43 ± 2·95 | 28 | 19·65 ± 2·91 | 26 | 19·02 ± 2·63 | 46 |
| **17** | **Minnesota** | Males | 16·13 ± 2·53 | 8 | 15·67 ± 1·97 | 6 | 15·65 ± 2·23 | 17 |
|  |  | Females | 15·62 ± 1·76 | 13 | 15·46 ± 2·11 | 13 | 15·26 ± 1·71 | 53 |
| **18** | **MPIP** | Males | 47·83 ± 13·06 | 47 | 48·07 ± 12·86 | 43 | 47·82 ± 12·80 | 154 |
|  |  | Females | 49·40 ± 12·78 | 65 | 49·60 ± 13·11 | 63 | 47·04 ± 13·51 | 200 |
| **19** | **Münster Neuroimaging Cohort** | Males | 35·27 ± 11·48 | 165 | 35·41 ± 11·86 | 163 | 37·22 ± 11·22 | 121 |
|  |  | Females | 34·94 ± 12·67 | 213 | 35·26 ± 12·15 | 211 | 38·27 ± 12·47 | 164 |
| **20** | **NESDA** | Males | 37·43 ± 9·66 | 7 | 34·50 ± 9·33 | 4 | 38·80 ± 10·81 | 15 |
|  |  | Females | 39·84 ± 9·74 | 19 | 39·31 ± 10·06 | 16 | 36 ·92 ± 10·07 | 72 |
| **21** | **Novosibirsk** | Males | NA | NA | NA | NA | NA | NA |
|  |  | Females | 41·57 ± 11·07 | 7 | 47·40 ± 6·99 | 5 | 43·71 ± 12·66 | 14 |
| **22** | **Oxford** | Males | 33·88 ± 9·19 | 8 | 32·00 ± 13·62 | 5 | 27·86 ± 10·10 | 14 |
|  |  | Females | 28·91 ± 10·89 | 11 | 27·14 ± 6·94 | 7 | 31·46 ± 10·80 | 24 |
| **23** | **QTIM** | Males | 22·30 ± 2·89 | 60 | 21·73 ± 3·64 | 56 | 21·73 ± 3·10 | 11 |
|  |  | Females | 22·53 ± 3·11 | 107 | 21·97 ± 3·51 | 104 | 22·20 ± 3·52 | 46 |
| **24** | **San Francisco UCSF** | Males | 15·16 ± 1·18 | 25 | 15·10 ± 1·18 | 21 | 15·15 ± 1·35 | 26 |
|  |  | Females | 15·59 ± 1·37 | 22 | 15·45 ± 1·43 | 20 | 15·88 ± 1·30 | 48 |
| **25** | **Sao Paulo (Wellcome)** | Males | 27·88 ± 7·23 | 25 | 28·43 ± 7·06 | 23 | 29·86 ± 9·97 | 7 |
|  |  | Females | 32·70 ± 8·77 | 20 | 32·60 ± 8·96 | 20 | 28·59 ± 7·90 | 17 |
| **26** | **SHIP/TREND** | Males | 49·98 ± 13·59 | 249 | 50·08 ± 14·31 | 248 | 47·70 ± 9·57 | 103 |
|  |  | Females | 49·73 ± 13·36 | 199 | 49·86 ± 13·44 | 197 | 49·34 ± 12·40 | 200 |
| **27** | **SHIP** | Males | 54·32 ± 11·81 | 114 | 54·36 ± 11·91 | 111 | 52·06 ± 9·55 | 36 |
|  |  | Females | 53·76 ± 11·91 | 95 | 53·69 ± 11·23 | 93 | 52·89 ± 10·96 | 95 |
| **28** | **Stanford** | Males | 36·27 ± 13·10 | 11 | 38·13 ± 8·22 | 8 | 37·19 ± 10·03 | 21 |
|  |  | Females | 37·73 ± 10·64 | 15 | 38·25 ± 11·19 | 12 | 35·67 ± 10·16 | 24 |
| **29** | **Sydney** | Males | 45·30 ± 22·22 | 20 | 46·15 ± 22·00 | 20 | 35·07 ± 21·25 | 68 |
|  |  | Females | 42·52 ± 22·52 | 29 | 43·71 ± 23·06 | 28 | 33·54 ± 20·36 | 137 |
|  | **Total** |  |  | **N=2,533** |  | **N=2,415** |  | **N=3,211** |

***Supplementary Table S1:*** **ENIGMA - Major Depressive Disorder Working Group Demographics.** Age (in years), and MDD patients-control breakdown for participating sites, separately for males and females, and training and test samples.

| **Cohort** | | **Sex** | **% Antidepressant users** | **% First episode MDD/Recurrent episode MDD** | **% Acute MDD/ Remitted MDD** | **Age of onset MDD (mean ± SD)** | **HDRS-17**^a^ **Severity MDD (mean ± SD)** | **BDI-II**^b^ **Severity MDD (mean ± SD)** |
| --- | --- | --- | --- | --- | --- | --- | --- | --- |
| **1** | **Barcelona** | Males | NA | NA | NA | NA | NA | NA |
|  |  | Females | 96 | 31/69 | 60/40 | 32·4 ± 11·6 | 12·8 ± 8·8 | NA |
| **2** | **BiDirect** | Males | 0 | 54/46 | 100/0 | 39·5 ± 10·0 | 12·5 ± 6·8 | NA |
|  |  | Females | 0 | 58/42 | 100/0 | 37·8 ± 11·3 | 14·3 ± 6·6 | NA |
| **3** | **BRDECC London** | Males | 77 | 0/100 | NA | 17·6 ± 5·7 | NA | 17·1 ± 13·4 |
|  |  | Females | 70 | 0/100 | NA | 21·9 ± 10·4 | NA | 17·5 ± 11·5 |
| **4** | **Edinburgh (Bipolar Family Study)** | Males | 0 | NA | NA | 20·2 ± 3·9 | 2·0 ± 3·2 | NA |
|  |  | Females | 27 | NA | NA | 22·5 ± 2·8 | 7·2 ± 6·9 | NA |
| **5** | **Calgary** | Males | 63 | 0/100 | 100/0 | 15·1 ± 2·0 | 15·3 ± 7·7 | 24·5 ± 10·5 |
|  |  | Females | 60 | 0/100 | 100/0 | 13·3 ± 1·9 | 16·2 ± 7·9 | 26·4 ± 14·4 |
| **6** | **CLiNG** | Males | 100 | 52/43 | 91/9 | 34·5 ± 10·0 | 20·5 ± 4·5 | 20·6 ± 14·5 |
|  |  | Females | 88 | 38/62 | 96/4 | 26·7 ± 9·8 | 19·3 ± 4·1 | 22·6 ± 6·4 |
| **7** | **CODE (Berlin)** | Males | 0 | 0/100 | 100/0 | NA | NA | NA |
|  |  | Females | 0 | 0/100 | 100/0 | NA | NA | NA |
| **8** | **Dublin** | Males | 77 | 31/69 | 100/0 | 29·4 ± 10·8 | 23·6 ± 4·5 | 21·4 ± 5·5 |
|  |  | Females | 83 | 27/73 | 100/0 | 26·3 ± 11·1 | 23·5 ± 5·4 | 19·8 ± 4·4 |
| **9** | **EPISCA (Leiden)** | Males | NA | NA | NA | NA | NA | NA |
|  |  | Females | 13 | 100/0 | 100/0 | NA | NA | NA |
| **10** | **FOR2107 - Marburg** | Males | 58 | 25/75 | 84/16 | 26·8 ± 13·7 | 8·4 ± 6·1 | 20·1 ± 10·2 |
|  |  | Females | 65 | 29/71 | 75/25 | 26·8 ± 12·6 | 8·0 ± 6·6 | 18·2 ± 11·5 |
| **11** | **FOR2107 - Münster** | Males | 81 | 45/55 | 76/24 | 25·9 ± 10·1 | 8·9 ± 7·2 | 18·3 ± 12·3 |
|  |  | Females | 50 | 35/65 | 69/31 | 25·1 ± 11·5 | 9·4 ± 7·6 | 15·8 ± 11·6 |
| **12** | **Groningen (DIP)** | Males | NA | NA | NA | NA | NA | NA |
|  |  | Females | 36 | 25/75 | 100/0 | 26·4 ± 16·0 | NA | NA |
| **13** | **Houston** | Males | 0 | 29/71 | 100/0 | 22·7 ± 9·8 | 12·0 ± 8·5 | 14·8 ± 13·8 |
|  |  | Females | 0 | 23/77 | 95/5 | 21·1 ± 11·1 | 9·7 ± 7·7 | 17·5 ± 15·6 |
| **14** | **Houston adolescents** | Males | 60 | 80/20 | 73/27 | 10·3 ± 2·3 | 8·3 ± 5·6 | NA |
|  |  | Females | 10 | 56/44 | 100/0 | 11·1 ± 2·0 | 14·7 ± 6·5 | NA |
| **15** | **McMaster University Mood Disorders** | Males | 50 | 50/50 | 100/0 | 20·8 ± 12·2 | 12·3 ± 8·5 | NA |
|  |  | Females | 63 | 37/63 | 100/0 | 24·1 ± 9·8 | 11·5 ± 6·8 | NA |
| **16** | **Melbourne** | Males | 29 | 35/65 | 100/0 | 16·7 ± 2·8 | NA | NA |
|  |  | Females | 26 | 36/64 | 100/0 | 16·4 ± 2·9 | NA | NA |
| **17** | **Minnesota** | Males | 19 | 10/90 | 0/100 | 12·5 ± 2·6 | NA | 20·8 ± 11·5 |
|  |  | Females | 25 | 54/46 | 0/100 | 12·4 ± 2·3 | NA | 27·8 ± 11·9 |
| **18** | **MPIP** | Males | 84 | 31/69 | 87/13 | 35·9 ± 13·9 | 26·7 ± 6·4 | 13·6 ± 9·4 |
|  |  | Females | 83 | 26/74 | 86/14 | 34·3 ± 13·7 | 26·4 ± 8·0 | 14·5 ± 11·9 |
| **19** | **Münster Neuroimaging Cohort** | Males | 93 | 25/75 | 91/9 | 29·4 ± 12·0 | 21·2 ± 6·8 | 23·3 ± 9·5 |
|  |  | Females | 91 | 22/78 | 93/7 | 29·6 ± 11·9 | 21·8 ± 7·7 | 26·4 ± 11·1 |
| **20** | **NESDA** | Males | 47 | 47/53 | 100/0 | 29·6 ± 12·4 | NA | NA |
|  |  | Females | 35 | 44/56 | 100/0 | 24·5 ± 11·0 | NA | NA |
| **21** | **Novosibirsk** | Males | NA | NA | NA | NA | NA | NA |
|  |  | Females | 71 | 14/86 | 64/36 | 36·4 ± 11·9 | 27·0 ± 3·9 | NA |
| **22** | **Oxford** | Males | 0 | 50/50 | 100/0 | 24·1 ± 10·5 | 22·7 ± 5·7 | NA |
|  |  | Females | 0 | 50/50 | 100/0 | 26·5 ± 8·3 | 23·1 ± 3·3 | NA |
| **23** | **QTIM** | Males | 18 | NA | NA | 17·5 ± 4·5 | NA | NA |
|  |  | Females | 22 | NA | NA | 18·7 ± 3·4 | NA | NA |
| **24** | **San Francisco UCSF** | Males | 0 | 48/52 | 96/4 | 13·1 ± 2·2 | NA | 26·0 ± 11·9 |
|  |  | Females | 0 | 29/71 | 88/12 | 13·3 ± 2·4 | NA | 27·1 ± 12·0 |
| **25** | **Sao Paulo (Wellcome)** | Males | 71 | 25/75 | 100/0 | NA | 15·0 ± 10·3 | NA |
|  |  | Females | 47 | 33/67 | 100/0 | NA | 15·6 ± 9·5 | NA |
| **26** | **SHIP/TREND** | Males | 14 | 44/56 | NA | 36·2 ± 12·8 | NA | 12·1 ± 8·4 |
|  |  | Females | 20 | 33/67 | NA | 36·0 ± 14·4 | NA | 12·5 ± 8·0 |
| **27** | **SHIP** | Males | 22 | 58/42 | NA | 39·0 ± 12·9 | NA | 11·8 ± 9·4 |
|  |  | Females | 17 | 55/45 | NA | 37·7 ± 12·8 | NA | 12·0 ± 10·8 |
| **28** | **Stanford** | Males | 44 | 10/90 | 100/0 | 20·2 ± 9·7 | NA | 30·1 ± 10·6 |
|  |  | Females | 45 | 18/82 | 100/0 | 18·8 ± 10·5 | NA | 21·9 ± 9·7 |
| **29** | **Sydney** | Males | 46 | 34/66 | 11/89 | 25·6 ± 18·8 | 11·3 ± 6·6 | NA |
|  |  | Females | 67 | 24/76 | 23/77 | 21·5 ± 13·2 | 13·1 ± 7·1 | NA |

***Supplementary Table S2:* ENIGMA - Major Depressive Disorder Working Group Clinical characteristics of MDD patients.** Percentage of MDD patients using antidepressant medication, percentage of first episode and recurrent episode MDD patients, percentage of acutely depressed and remitted MDD patients, age of onset of MDD and severity of symptoms breakdown for participating sites, separately for male and female samples.

^a^ Measured with the Hamilton Depression Rating Scale (HDRS-17; range: 0-52)

^b^ Measured with the Beck Depression Inventory (BDI-II; range: 0-63)

| **Cohort** | **Country** | **Scanner type** | **Sequence T1** | **FreeSurfer version** | **Slice orientation** | **Operating system** |
| --- | --- | --- | --- | --- | --- | --- |
| **Barcelona** | **Spain** | 3T Philips Achieva | 3D MPRAGE images (Whole-brain T1-weighted); TR=6.7ms, TE=3.2ms; 170 slices, voxel size 0.89X0.89X1.2 mm. Image dimensions 288X288X170; field of view: 256X256X204; slice thickness: 1.2 mm; with a sagittal slice orientation, T1 contrast enhancement, flip angle: 8º, grey matter as a reference tissue, ACQ matrix MXP = 256X240 and turbo-field echo shots (TFE) = 218. | 6 | Sagittal | Scientific Linux 5 |
| **BiDirect** | **Germany** | 3 T Philips Intera scanner | 3D T1-weighted turbo field echo images were collected with a the following parameters: TR = 7.26, TE = 3.56, 9° flip angle, 160 sagittal slices, matrix dimension 256 x 256, FOV = 256 x 256mm, 2mm slice thickness (reconstructed to 1mm) and a resulting voxel size of 1x1x1mm | 5.3 | Sagittal |  |
| **Edinburgh (Bipolar Family Study)** | **Scotland** | 1.5T GE Signa | T1-weighted sequence. TR=500 msec; TE=4 msec; flip angle 8°; matrix 192 x 192; 180 slices; voxel size 1.25 mm x 1.25 mm x 1.20 mm; FOV=24, phase FOV 1 | 5.3 | Coronal | linux 6, x86_64, kernel 2.6.32 |
| **BRCDECC London** | **England** | 1.5T GE Signa HDx | ADNI-1 MPRAGE pulse sequence (details at http://adni.loni.ucla.edu/research/protocols/mri-protocols/) | 5.3 | Sagittal | Linux-centos4_x86_64 |
| **Calgary** | **Canada** | 1.5T Siemens Magnetom Vision. 3T GE Discovery MR750 | 1.5T: A sagittal scout series was acquired to test image quality. 3D fast low angle shot (FLASH) sequence was used to acquire data from 124 1.5 mm-thick contiguous coronal slices through the entire brain (echo time = 5ms, repetition time = 25ms, acquisition matrix = 256 x 256 pixels, field of view = 24 cm and flip angle = 40°). 3T: Anatomical imaging acquisition parameters: axial acquisition, repetition time (TR), 2200 milliseconds (ms); echo time (TE), 3.04 ms; TI, 766, 780; flip angle, 13 degrees; 208 partitions; 256 × 256 matrix; and field of view, 256. | 5.3 | Dalhousie sample, coronal; Calgary sample, axial | MacOs Sierra |
| **CliNG** | **Germany** | 3T Siemens Tim Trio | T1-weighted 3D MPRAGE; TR/TE/TI/FA=2250 ms/3.26 ms/900 ms/9°; image matrix = 256 x 256; 192 sagittal slices; voxel size= 1 mm3 | 5.3 | Sagittal | Linux |
| **CODE (Berlin)** | **Germany** | 3T Siemens Trio (4 Sites), 3 T Philips Achieva (1 site) | Siemens: T1 mprage, voxel size 1 mm x 1 mm x 1 mm; TR=1900 msec; TE=2.52 msec; Sample 1: 192 slices, Sample 2: 176 slices (except 1 site: 192) Philips: T1 3D-TFE, voxel size 1 mm x 1 mm x 1 mm; TR=8.3 msec; TE=3.8 msec; 170 slices. | 5.3 | Sagittal | Ubuntu 12.04 LTS (Linux 64bit) |
| **Dublin** | **Ireland** | 3T Phillips Achieva; 1.5T Siemens Vision | 3T: A sagittal T1 3D TFE was used to scan all participants. TR=8.5 msec; TE=3.9 msec; FOV = 256 mm, AP: 256 mm, RL: 160 mm; matrix: 256×256. 1.5T: 3D-MPRAGE T1-weighted sequence. TR=11.6 msec; TE=4.9 msec; FOV=230 mm; matrix 512 x 512, slice thickness: 1.5 mm. | 5.3 | Sagittal (3T), Coronal (1.5T) | Mac OS |
| **EPISCA (Leiden)** | **The Netherlands** | 3T Philips Achieva | a sagittal 3-dimensional gradient-echo T1-weighted image was acquired (repetition time = 9.8 ms; echo time = 4.6 ms; flip angle = 8°; 140 sagittal slices; no slice gap; field of view =256 × 256 mm; 1.17 × 1.17 × 1.2 mm voxels; duration = 4:56 min) | 5.3 | Sagittal | Ubuntu 14.04.5 LTS (Linux 3.13.0-153-generic x86_64) |
| **FOR2107 - Marburg** | **Germany** | 3T Siemens Magnetom TiroTim syngo MR B17 | Sequence: 3D T1-weighted magnetization prepared rapid acquisition gradient echo (MPRAGE) - Sagittal Acquisition Direction, # of Slices 176, 0.5mm Slice Gap, 1.0x1.0x1.0 Voxel Size (mm3), TI 900 ms, TE 2.26 ms, TR 1900 ms, Flip Angle 9. | 5.3 | Sagittal | Red Hat Enterprise Linux Server release 5.11 (Tikanga) |
| **FOR2017 - Münster** | **Germany** | 3T Siemens PRISMA | Sequence: 3D T1-weighted magnetization prepared rapid acquisition gradient echo (MPRAGE). - Sagittal Acquisition Direction, # of Slices 192, 0mm Slice Gap, 1.0x1.0x1.0 Voxel Size (mm3), TI 900 ms, TE 2.28 ms, TR 1900 ms, Flip Angle 8 | 5.3 | Sagittal | Red Hat Enterprise Linux Server release 5.11 (Tikanga) |
| **Groningen sample (DIP)** | **The Netherlands** | 3T Philips | 3D T1-weighted scan (170 slices; TR = 9ms; TE = 3.6ms; 256x231 matrix of 1×1×1 mm voxels) | 5.3 |  |  |
| **Houston** | **USA** | subjects in 20000s: 1.5 T Philips Medical Systems Gyroscan Intera; subjects in 30000s: 3T Siemens Allegra | Subjects in the 20000s: Fast field echo sequence- repetition time (TR) = 24 ms, echo time (TE) = 4.99 ms, flip angle = 40°, slice thickness = 1 mm, matrix size = 256 × 256 and 150 slices. Subjects in 30000s: MPRAGE- repetition time (TR) = 1750 ms, echo time (TE) = 4.39 ms, flip angle = 8°, slice thickness = 1 mm, matrix size = 208 × 256 and 160 slices. | 5.3 | Subjects in 20000s: Sagittal; Subjects in 30000s: Transverse | Fedora 19 |
| **Houston adolescents** | **USA** | subjects in 20000s: 1.5 T Philips Medical Systems Gyroscan Intera; subjects in 30000s: 3T Siemens Allegra | Subjects in the 20000s: Fast field echo sequence- repetition time (TR) = 24 ms, echo time (TE) = 4.99 ms, flip angle = 40°, slice thickness = 1 mm, matrix size = 256 × 256 and 150 slices. Subjects in 30000s: MPRAGE- repetition time (TR) = 1750 ms, echo time (TE) = 4.39 ms, flip angle = 8°, slice thickness = 1 mm, matrix size = 208 × 256 and 160 slices. | 5.3 | Subjects in 20000s: Sagittal; Subjects in 30000s: Transverse | Fedora 19 |
| **McMaster University Mood Disorders** | **Canada** | 1.5T (GE); 3T(GE) | 1.5-T. Sigma GE Genesis-based Echo-Speed scanner running version 5.7 software and using a standard 30-cm circularly polarized head coil. Sagittal anatomic images were acquired by using a 3D/FSPGR/20 sequence (flip angle=20; echo delay time in-phase (TE), minimum repetition time (TR)=300 ms; inversion recovery=300 ms; matrix=512x256; field of view (FOV)=24 cm; scan thickness=1.2 mm). 3-T MRI Sigma GE Genesis (General Electric Medical Systems, Milwaukee, WI). Sagittal T-1 weighted images were acquired using a 3D FSPGR-IR sequence, (TR/TE=10.3/2.1 ms; flip angle=20; inversion time=300; matrix=512x256; FOV=24; and slice thickness=1.2 mm. | 5 |  |  |
| **Melbourne** | **Australia** | 3T GE Signa Excite | 3D BRAVO sequence 140; TR=7900 ms; TE=3000 ms; flip angle=13º; FOV=256 mm; matrix=256 x 256 | 5.3 | Axial | Linux Debian x86 64 |
| **Minnesota** | **USA** | 3.0 Tesla Tim Trio scanner; Siemens Corp | A 5-minute structural scan was acquired using a T1-weighted, high-resolution, magnetization-prepared gradient-echo sequence: repetition time, 2530 milliseconds; echo time, 3.65 milliseconds; inversion time, 1100 milliseconds; flip angle, 7°; field of view, 256 × 176 mm; voxel size, 1-mm isotropic; 224 slices; and generalized, autocalibrating, partially parallel acquisition acceleration factor, 2. | 5.3 | Coronal | Linux |
| **MPIP** | **Germany** | 1.5T GE and Siemens (the latter: only few cases) | #1: T1-weighted SPGR sagittal 3D volume. TR=1030 msec; TE=3.4 msec; 124 slices; matrix=256x256; FOV=23.0x23.0 cm2; voxel size=0.8975 mm x0.8975 mm x 1.2- 1.4 mm; flip angle=90°; birdcage resonator. #2: same scanner as #1, platform update Signa Excite, sagittal T1-weighted (spin echo sequence, TR=9.7 msec, TE=2.1 msec; FOV=25.0x25.0 cm2, voxel size=0.875 mm x0.875 mm x1.2 mm, 124- 132 slices, flip angle=90°. #3: Siemens 1.5 Tesla, Vario, 3D MPRAGE, TR=11.6 msec; TE=4.9 msec; FOV 23x23 cm2; matrix 512x512; 126 axial slices; voxel site 0.45 mm x 0.45 mm x 1.5 mm. (only N=2 subjects) | 5.3 | 1.5 GE: sagittal. 1.5 Siemens: axial | Linux 2.6.37.1-1.2- desktop x86_64 |
| **Münster Neuroimaging Cohort** | **Germany** | 3T Philips Gyroscan Intera | 3D fast gradient echo sequence (turbo field echo), repetition time = 7.4 milliseconds, echo time = 3.4 milliseconds, flip angle = 9°, two signal averages, inversion prepulse every 814.5 milliseconds, acquired over a field of view of 256 (feet -head [FH]) × 204 (anterior -posterior [AP]) × 160 (right -left [RL]) mm, phase encoding in AP and RL direction, reconstructed to cubic voxels of .5 mm × .5 mm × .5 mm | 5.3 | Sagittal | Red Hat Enterprise Linux Server release 5.11 (Tikanga) |
| **NESDA** | **The Netherlands** | 3T Phillips Achieva/Intera | 3D gradient-echo T1-weighted sequence. TR=9 msec; TE=3.5 msec; flip angle 8º, FOV = 256 mm; matrix: 25x62x56; in plane voxel size = 1 mm × 1 mm x 1 mm; 170 slices. | 5 | Sagittal | SHARK HPC, Linux environment |
| **Novosibirsk** | **Russia** | 3T GE Discovery™ MR750w | Whole-brain T1-weighted images - 3D fast spin gradient echo sequence (FSPGR BRAVO), repetition time = 9.5 ms, echo time = 3.7 ms, flip angle = 3°, acquired over a field of view of 256 (feet-head [FH]) × 256 (anterior-posterior [AP]) × 188 (rightleft [RL]) mm, reconstructed to cubic voxels of 1 mm × 1 mm × 1 mm | 5.3 | Sagittal | OS X 10.10 |
| **Oxford** | **England** | 3T Siemens Tim Trio | Voxel resolution 0.78 x 0.8 x 0.78 mm on a 208 x 256 x 200 grid, TE/TI/TR= 4.8/1100/2040 ms | 5.3 |  |  |
| **QTIM** | **Australia** | Bruker 4T Wholebody MRI | 3D T1 weighted sequence. TR=1500 msec; TE=3.35 msec; flip angle=8°, 256 or 240 (coronal or sagittal) slices, FOV=240 mm, matrix 256x256x256 (or 256x256x240) | 5.1 | Coronal, then sagittal following software upgrade. | Linux- centos4_x86_64- stable-pub-v5.1.0 |
| **San Francisco UCSF** | **USA** | 3T GE Discovery MR750 | SPGR T1-weighted: TR=8.1 ms; TE=3.17 ms; TI=450 ms; flip angle=12°; 256x256 matrix; FOV=250x250 mm; 168 sagittal slices; slice thickness=1 mm; in-plane resolution=0.98x 0.98 mm | 5.3 | Sagittal | Linux-centos6_x86_64-stable-pub-v5.3.0. |
| **Sao Paulo (Wellcome)** | **Brasil** | 1.5T General Eletric (GE) | Imaging data were acquired using two MRI scanners (at the Clinics Hospital of the University of Sa ̃ o Paulo 1.5 T GE Signa scanner, General Electric, Milwaukee Wisconsin, USA). T1-SPGR sequence providing 124 contiguous slices, voxel size 0.8660.8661.5 mm, echo time 5.2 ms, resolution time 21.7 ms, flip angle 20, field of vision 22, matrix 256x192) | 5.3 |  |  |
| **SHIP** | **Germany** | 1.5T Siemens Avanto | 3D T1-weighted (MP-RAGE/ axial plane); TR=1900 msec; TE=3.4 msec; Flip angle=15°; voxel size 1 mm x 1 mm x 1 mm | 5.3 (cortical), 5.1 (subcortical) | Axial | Centos6_x86_64 |
| **SHIP/TREND** | **Germany** | 1.5T Siemens Avanto | 3D T1-weighted (MP-RAGE/ axial plane); TR=1900 msec; TE=3.4 msec; Flip angle=15°; voxel size 1 mm x 1 mm x 1 mm | 5.3 (cortical), 5.1 (subcortical) | Axial | Centos6_x86_64 |
| **Stanford** | **USA** | 1.5T GE Signa Excite | Whole-brain T1-weighted images were collected using a spoiled gradient echo (SPGR) pulse sequence (116 sagittal slices; through-plane resolution = 1.5 mm; in-plane resolution = 0.86 x 0.86 mm; flip angle = 15 degrees; repetition time [TR] = 8.3-10.1 ms; echo time [TE] = 1.7-3.0; inversion time [TI] = 300 ms; matrix = 256 x 192). | 5.3 | Sagittal | Linux-centos6_x86_64 |
| **Sydney** | **Australia** | 3T GE MR750 | 3D T1-weighted sequence. TR=7.2 msec; TE=2.78 msec; matrix =256; FOV=240; No. slices=196; thick=0.9mm; inplane resolution=0.9375 | 5.1 but rerunning it for 5.3 | Coronal | Linux_Ubuntu16.04 lts 64bit |

***Supplementary Table S3*: ENIGMA - Major Depressive Disorder Working Group Image acquisition and processing by site.**

| **Cohort** | **Country** | **Diagnosis measurment** | **Sample characteristics/Inclusion criteria** | **Exclusion criteria** |
| --- | --- | --- | --- | --- |
| **Barcelona** | **Spain** | DSM-IV-TR acc. to CIDI-interview and HAMD | Outpatients with MDD diagnosis (DSM-IV-TR), with a first episode, recurrent MDD or chronic MDD (TRD) age 18-65 | The exclusion criteria for healthy participants were: lifetime psychiatric diagnoses, first-degree relatives with psychiatric diagnoses and clinically significant physical or neurological illnesses. Axis I comorbidity according to DSM-IV-TR criteria was an exclusion criteria for all participants. |
| **BiDirect** | **Germany** | M.I.N.I. Neuropsychiatric Interview, IDS, HAMD, CESD, ICD-10 | Patients hospitalized for a first or recurrent episode of depression, population controls randomly selected in city registry | dementia, addiction |
| **Edinburgh (Bipolar Family Study)** | **Scotland** | SCID interview | The MDD group were originally people with a FHx of bipolar disorder | MDD subjects: presence of other axis I diagnoses. Control subjects: no medical history, including neurological and psychiatric history, as well as no previous or actual use of psychotropic medication All subjects: any major neurological disorder, learning disability, or any history of head injury that included loss of consciousness and any contraindications to MRI. |
| **BRCDECC London** | **England** | SCAN interview | Community based or outpatients, none were inpatients. MDD subjects: Less than two depressive episodes of at least moderate severity. Did not meet DSM-IV diagnostic criteria for recurrent major depressive disorder. Control group participants were clinically interviewed to ensure they had never experienced depressive symptoms.  Exclusion criteria for all participants were for contraindications to MRI; other exclusion criteria were a diagnosis of neurological disorder, head injury leading to loss of consciousness or conditions known to affect brain structure or function (including alcohol or substance misuse), ascertained during clinical interview. Potential participants were also excluded if they or a first-degree relative had ever fulfilled criteria for mania, hypomania, schizophrenia or mood-incongruent psychosis. | Contraindications to MRI, diagnosis of neurological disorder, head injury leading to loss of consciousness or conditions known to affect brain structure or function (including alcohol or substance misuse), if they or a first-degree relative had ever fulfilled criteria for mania, hypomania, schizophrenia or mood-incongruent psychosis. |
| **Calgary** | **Canada** | KSADS | First episode MDD and healthy controls (Dalhousie sample). Recurrent MDD and healthy controls, recruited via referral from clinicians in Calgary, Alberta and through advertisements in local clinics and at the University of Calgary (Calgary sample). | Dalhousie Sample: A history of neurological illness, medical illness, claustrophobia, >21 year of age, or the presence of a ferrous implant or pacemaker. University of Calgary: Left handed; history of seizures, epilepsy or other neurological or psychiatric diagnoses (specifically bipolar disorder, psychosis, pervasive developmental disorder, eating disorders, PTSD); pregnancy |
| **CliNG** | **Germany** | ICD-10 interview | Patients met the diagnostic criteria for major depressive disorder according to ICD-10 classification standards and were aged between 18 and 60 years. | Exclusion criteria for MDD subjects were neurological and severe other medical conditions (in particular those that could be related to affective symptoms), lifetime diagnosis of substance dependence, substance abuse during the last month, cannabis abuse during the last 2 weeks, mental retardation as well as past or actual presence of other axis I diagnoses with exception of anxiety disorders. Exclusion criteria for control subjects were neurological, psychiatric and severe other medical conditions, lifetime diagnosis of substance dependence, substance abuse during the last month, cannabis abuse during the last 2 weeks, previous or actual use of psychotropic medication, and mental retardation. |
| **CODE (Berlin)** | **Germany** | SCID interview | Chronic depression, at least for 1 year, subjects medication free (CBASP1); Chronic depression, at least 2 years, early onset (before 21 years), medicaton free (CBASP2) | MDD: Presence of any other Axis-1 diagnosis; Acute risk for suicide (in contrast to suicidal ideation); History of psychotic symptoms, bipolar disorder, or dementia; Schizotypal, antisocial or borderline personality disorder; Use of psychotropic medication within two weeks prior to the start of the study; No current psychotherapeutic treatment. Control subjects: No history of or current Axis-1 or 2 disorders. All subjects: History of or current neurological disorder or brain injury; Serious medical condition; Severe cognitive impairment; Substance-related abuse or dependence disorder; Use of psychotropic medication; Use of central-acting medication; Pregnancy; General MRI contraindications. |
| **Dublin** | **Ireland** | SCID-1 interview |  | MDD subjects: comorbid psychiatric disorders (Axis I or Axis II, other than MDD), Treatment with antipsychotics or mood stabilizers, age 65, Control subjects: no Axis-I diagnosis, no medication use. All subjects: history of neurological or other severe medical illness, head injury or severe substance abuse in their lifetime history and general MRI contraindications. |
| **EPISCA (Leiden)** | **The Netherlands** | ADIS | Inclusion criteria for the patient group were: having clinical depression  as assessed by categorical and dimensional measures of DSM-IV depressive and anxiety disorders, no current and prior use of antidepressants, and being referred for CBT at an outpatient care unit. Inclusion criteria for the control group were:  no current or past DSM-IV classifications, no clinical scores on validated  mood and behavioral questionnaires, no history of traumatic experiences, and no current psychotherapeutic and/or psychopharmacological intervention of any kind. | Primary DSM-IV clinical diagnosis of ADHD, ODD, CD, pervasive developmental disorders, post-traumatic stress disorder, Tourette's syndrome, obsessive–compulsive disorder, bipolar disorder, and psychotic disorders; current substance abuse; history of neurological disorders or severe head injury; age < 12 or > 21 years; pregnancy; left-handedness; IQ score < 80 as measured by the Wechsler Intelligence Scale for Children (WISC) (Wechsler, 1991) or Adults (Wechsler, 1997); and general MRI contra- indications. |
| **FOR2107 - Marburg** | **Germany** | SCID-1 | Participants recruited by means of public advertisement and from the inpatient services. Inclusion criteria: age 18-65 years; patients were diagnosed with major depressive disorder by SCID-Interview, currently depressed or remitted. | Exclusion criteria all: any MRI contraindications; any neurological abnormalities. Exclusion criteria controls: any current or former psychiatric disorder; Exclusion criteria patients: substance dependence or current benzodiazepine treatment (wash out of at least three half-lives before study participation)" |
| **FOR2017 - Münster** | **Germany** | SCID-1 | Participants recruited by means of public advertisement and from the inpatient services. Inclusion criteria: age 18-65 years; patients were diagnosed with major depressive disorder by SCID-Interview, currently depressed or remitted. | Exclusion criteria all: any MRI contraindications; any neurological abnormalities. Exclusion criteria controls: any current or former psychiatric disorder; Exclusion criteria patients: substance dependence or current benzodiazepine treatment (wash out of at least three half-lives before study participation)" |
| **Groningen sample (DIP)** | **The Netherlands** | MINI-SCAN | Outpatients with MDD diagnosis. Inclusion MDD: Outpatients treated in mental health care for depression, BDI-II>13 at screening, adults. | Exclusion MDD: Comorbid axis-I disorders other than anxiety disorders or past substance abuse, other psychotropic medication than stable use of SSRI/SNRI/TCA, established cardiovascular disease, active and concrete suicidal plans, inadequate language proficiency, cognitive impairments or neurological disease that interferes with task performance. Exclusion CTL: Same as MDD, lifetime history of MDD, BDI>8. |
| **Houston** | **USA** | SCID interview | Outpatients | MDD subjects: age below 18; lifetime or current diagnosis of psychotic disorder, or bipolar I or II disorder; substance abuse/dependence in 6 months prior to study inclusion; current major medical problems. Control subjects: age below 18; current major medical problems; current psychiatric or neurologic disorder; history of psychiatric disorders in a first-degree relative; current major medical problems. Both groups: MRI contra-indications |
| **Houston adolescents** | **USA** | Major depressive disorder (MDD) diagnosis according to DSM-IV | Outpatients | MDD subjects: head trauma with residual effects, neurological disorders, uncontrolled major medical conditions based on patient self reports and current drug abuse. In addition, Healthy controls (HC) were excluded if they had a history of any Axis I disorder or had a first-degree relative with any Axis I disorder. |
| **McMaster University Mood Disorders** | **Canada** | SCID | Outpatients | Comorbid Axis 1 disorders excluded, including for example, psychosis, bipolar, PTSD substance dependence or current active eating disorder. Exclusion criteria included: i) treatment with anti-cholinergic or typical (first generation) anti-psychotic medication; ii) electroconvulsive therapy (ECT) or transcranial magnetic stimulation (TMS) within the past year; iii) a history of substance dependence or significant and recent (< 1 year) substance abuse; iv) a history (within the past 12 months) of an endocrine or other medical disorder known to adversely affect cognition (e.g., Cushing’s, uncontrolled diabetes, seizure disorder); and v) English comprehension lower than a grade 6 reading level. |
| **Melbourne** | **Australia** | SCID interview | Youth depression sample: 15-25 years of age. Recruited as part of 2 large RCTs (incl. YoDA-C - Davey et al., 2014; Trials) and scanned prior to treatment randomisation. 60 patients unmedicated (YoDA-C). | MDD subjects: lifetime or current SCID-I diagnosis of psychotic disorder, or bipolar I or II disorder. Control subjects: any SCID-I diagnosis or medication use. Both groups: Acute or unstable medical disorder; general MRI contraindications |
| **Minnesota** | **USA** | Schedule for Affective Disorders and Schizophrenia for School-Age Children–Present and Lifetime Version and the Children’s Depression Rating Scale–Revised (CDRS-R). | Adolescents with MDD and HCs aged 12 to 19 years were recruited to participate through community postings and referrals from local mental health services. Adolescents with MDD were eligible if they had a primary diagnosis of MDD and had not received any psychotropic medication treatment for the past 2 months. Healthy adolescents were eligible if they had no current or past psychiatric diagnoses and were frequency matched to the MDD group on age and sex | Exclusion criteria for both groups included the presence of a neurologic or other chronic medical condition, mental retardation, pervasive developmental disorder, substance use disorder, bipolar disorder, or schizophrenia |
| **MPIP** | **Germany** | M-CIDI/SCAN interview | M. A. R. S. sample: both first and recurrent episodes; RUD sample: only recurrent episodes with some patients scanned in remission | Munich Antidepressant Response Signature (MARS) study MDD subjects (clinical consensus diagnosis or M-CIDI (since 2008)): depressive syndromes secondary to any medical or neurological condition (e. g., intoxication, drug abuse, stroke), the presence of manic, hypomanic or mixed affective symptoms, lifetime diagnosis of alcohol dependence, illicit drug abuse or the presence of severe medical conditions (e.g., ischemic heart disease). Patients with bipolar depression were excluded for the current MR study. Control subjects: age > 65, MMSE<27, presence of severe somatic diseases or lifetime history of the following axis I disorders as assessed by the M-CIDI interview: alcohol dependence, drug abuse or dependence, possible psychotic disorder, mood disorder, anxiety disorder including OCD and PTSD, somatoform disorder, dissociative disorder NOS, and eating disorder 2. Recurrent unipolar depression (RUD) study: MDD subjects (SCAN interview): presence of manic episodes, mood incongruent psychotic symptoms, the presence of a lifetime diagnosis of intravenous drug abuse and depressive symptoms only secondary to alcohol or substance abuse or to medical illness or medication.Control subjects: presence of severe somatic diseases or life-time history of anxiety and affective disorders according to the Composite International Diagnostic-Screener (CIDI-S). All subjects: gross incidental MR findings such as territorial infarction, tumor, hydrocephalus, malformations and anatomical deviations (e.g. enlarged ventricles) that prevent appropriate image processing were additional exclusion criteria. 3. MR images of 9 additional controls acquired at the LMU, Munich, meeting equivalent criteria as the RUD control sample were included. |
| **Münster Neuroimaging Cohort** | **Germany** | SCID interview | Participants recruited by means of public advertisement and from the inpatient services. Inclusion criteria: age 16-65 years; patients were diagnosed with major depressive disorder by SCID-Interview | MDD subjects: presence of bipolar disorder, schizoaffective disorders and schizophrenia; substancerelated disorders or current benzodiazepine treatment (wash out of at least three half-lives before study participation), and former electroconvulsive therapy. Control subjects: any current or former psychiatric disorder. Both groups: any neurological abnormalities, MRI contra-indications |
| **NESDA** | **The Netherlands** | CIDI interview | DSM-4 based diagnosis of MDD (6 month recency), using CIDI interview. 93 (60%) MDD patients have a comorbid ANX diagnosis. Age range 18-65 | MDD subjects: presence of axis-I disorders other than MDD, panic disorder, social anxiety disorder, or generalized anxiety disorder and any use of psychotropic medication other than stable use of SSRIs or infrequent benzodiazepine use (i.e., equivalent to 2 doses of 10 mg of oxazepam 3 times per week or use within 48 hours prior to scanning). Control subjects: no Axis-I diagnosis, no medication use. All subjects: presence or history of major internal or neurological disorder, dependence on or recent abuse (past year) of alcohol and/or drugs, hypertension, and general MRI contraindications. |
| **Novosibirsk** | **Russia** | MINI, SCID, ICD-10 interviews | MDD Patients: 100% hospital-based, outpatients: 0%, general population: 0% | MDD subjects: Presence of axis-I disorders other than MDD, panic disorder, social anxiety disorder, or generalized anxiety disorder and any use of psychotropic medication other than stable use of SSRIs or infrequent benzodiazepine use; age 18 or below; alcohol or substance abuse/dependence within 6 months of study participation; current major medical problems. Control subjects: age over 65; any current or former psychiatric disorder. Both groups: MRI contra-indications. |
| **Oxford** | **England** | SCID interview | Possibly another 15-20 MDD scans from another study; in general no comorbidities - some high anxiety levels and history of panic attacks; drug free | MDD: psychosis or substance dependence (DSM-IV), clinically significant risk of suicidal behaviour, having contraindications to escitalopram treatment or being treated with psychotropic medication less than three weeks before the study (five weeks in the case of fluoxetine); HC: current or past history of Axis I disorder as defined by DSM-IV; Both groups: major somatic or neurological disorders, pregnancy or breast-feeding, contra-indications to MR imaging or concurrent medication which could alter emotional processing |
| **QTIM** | **Australia** | CIDI interview | Retrospective questionnaire about depression episodes combined with an MRI study. The best described MDD episode is defined as the worst one (according to individuals). We have up to 5 supplementary episodes (briefly) described. Sample composed of twins and relatives. Population-based sample | MDD subjects: presence of axis-I disorders other than MDD and anxiety disorders Control subjects: antidepressant use, psychiatric disorders All subjects: relatedness between subjects, left handedness, history of neurological or other severe medical illness, head injury or current or past diagnosis of substance abuse, use of cognition affecting medication and general MRI contraindications |
| **San Francisco UCSF** | **USA** | KSADS (semi-structured interview based on DSM) for MDD, DISC/DPS for HCL | Outpatient/community-based sample with DSM diagnosis, mostly antidepressant-naive and approximately 60% of MDD have comorbid anxiety disorders | Exclusion criteria for all participants included: 1) use of pharmacotherapeutics for treating psychiatric conditions within the past 6 months, 2) misuse of drugs within two months prior to MRI scanning; 3) two or more alcoholic drinks per week within the previous month (as assessed by the Customary Drinking and Drug Use Record; CDDR) (Brown et al, 1998); 4) a full scale IQ score of less than 75 (as assessed by the Wechsler Abbreviated Scale of Intelligence; WASI) (Wechsler, 1999); 5) contraindications for MRI including ferromagnetic implants and claustrophobia; 6) pregnancy or the possibility of pregnancy; 7) left-handedness; 8) prepubertal status (as assessed as Tanner stages of 1 or 2) (Tanner, 1962); 9) inability to understand and comply with procedures; 10) neurological disorder (including meningitis, migraine, or HIV); 11) head trauma; 12) learning disability; 13) serious health problems; and 14) complicated or premature birth (i.e., birth before 33 weeks of gestation). The MDD group was subject to the additional exclusion criterion of a primary psychiatric diagnosis other than MDD. The HCL group was subject to the additional exclusion criteria of: 1) history of mood or psychotic disorders in a first- or second-degree relative (as assessed by the Family Interview for Genetics; FIGS) (Maxwell, 1992); and 2) current or lifetime DSM-IV-TR Axis I psychiatric disorder. |
| **Sao Paulo (Wellcome)** | **Brasil** | Hamilton Rating Scale for Depression (HRSD) | Population-based study of incident (first-episode) psychosis in outpatient services. All subjects we provided were diagnosed with psychotic depression (and not schizophrenia, bipolar disorder or other psychotic diagnoses). | People with psychotic disorders due to a general medical condition or substance-induced psychosis were excluded. Additional exclusion criteria were: (a) history of head injury; (b) presence of neurological disorders or any organic disorders that could affect the central nervous system; and (c) contraindications for MRI. Exclusion cri- teria specific for the control group were personal history of psychosis or other Axis I disorders, except substance misuse or mild anxiety disorders. |
| **SHIP** | **Germany** | M-CIDI interview | Population based longitudinal cohort study | MDD subjects: presence of axis-I disorders other than MDD, anxiety disorders, conversion, somatization and eating disorder. Control subjects: no lifetime diagnosis of depression, no antidepressiva, and severity index=0 All subjects: We removed subjects with medical conditions (e.g. a history of cerebral tumor, stroke, Parkinson’s diseases, multiple sclerosis, epilepsy, hydrocephalus, enlarged ventricles, pathological lesions) or due to technical reasons (e.g. severe movement artifacts or inhomogeneity of the magnetic field). |
| **SHIP/TREND** | **Germany** | M-CIDI interview | Population based longitudinal cohort study | MDD subjects: no special exclusion criteria Control subjects: no lifetime diagnosis of depression, no antidepressiva, and severity index=0 All subjects: We removed subjects with due to medical conditions (e.g. a history of cerebral tumor, stroke, Parkinson’s diseases, multiple sclerosis, epilepsy, hydrocephalus, enlarged ventricles, pathological lesions) or due to technical reasons (e.g. severe movement artifacts or inhomogeneity of the magnetic field). |
| **Stanford** | **USA** | SCID interview | Community-based DSM-diagnosed sample | MDD subjects: presence of axis-I disorders other than MDD, anxiety and eating disorders . Control subjects: control individuals did not meet diagnostic criteria for any current psychiatric. Both groups: alcohol / substance abuse or dependence within six months prior to MRI scanning, history of head trauma with loss of consciousness > 5 min, aneurysm, or any neurological or metabolic disorders that require ongoing medication or that may affect the central nervous system (including thyroid disease, diabetes, epilepsy or other seizures, or multiple sclerosis), MRI contraindications, or bad MRI data (e.g., extreme movement). |
| **Sydney** | **Australia** | SCID interview |  | MDD subjects: presence of axis-I disorders other than MDD, panic disorder, social anxiety disorder, or generalized anxiety disorder. Control subjects: no Axis-I diagnosis, no medication use. Exclusion criteria for all subjects included medical instability (as determined by a psychiatrist), history of neurological disease (e.g. tumour, head trauma, epilepsy), medical illness known to impact cognitive and brain function (e.g. cancer), intellectual and/or developmental disability and insufficient English for neuropsychological assessment. All subjects were asked to abstain from drug or alcohol use for 48 hours prior to testing and informed about a drug screen protocol. |

***Supplementary Table S4*: ENIGMA - Major Depressive Disorder Working Group Instrument for diagnosing Major Depressive Disorder and exclusion criteria by site.**

| **Machine learning algorithm** | **R** | | **R^2^** | | **MAE** | |
| --- | --- | --- | --- | --- | --- | --- |
|  | Male training sample | Female training sample | Male training sample | Female training sample | Male training sample | Female training sample |
| **Ridge regression** | 0·85 | 0·84 | 0·73 | 0·71 | 6·75 | 6·86 |
| **GPR linear** | 0·86 | 0·85 | 0·74 | 0·72 | 6·63 | 6·76 |
| **GPR RBF** | 0·87 | 0·87 | 0·75 | 0·76 | 6·35 | 6·14 |
| **GAM** | 0·87 | 0·86 | 0·75 | 0·74 | 6·45 | 6·56 |
| **SVR linear** | 0·85 | 0·84 | 0·72 | 0·71 | 6·86 | 6·91 |
| **SVR RBF** | 0·85 | 0·87 | 0·73 | 0·75 | 6·50 | 6·09 |

***Supplementary Table S5:* Alternative machines and kernels in the brain age prediction framework.** Performance metrics in the training samples of males and females across four different machine learning algorithms/kernels are displayed here. R, Pearson’s correlation; R2, explained variance; MAE, mean absolute error.

| **Age group** | **Brain-PAD** | | **MAE** | |
| --- | --- | --- | --- | --- |
|  | **Male test samples**  **(N=2,256)** | **Female test samples**  **(N=3,370)** | **Male test samples**  **(N=2,256)** | **Female test samples**  **(N=3,370)** |
| **10-19 years** | 4·29 (8·52) | 3·77 (8·08) | 7·19 (6·25) | 6·86 (5·69) |
| **20-29 years** | 3·59 (7·35) | 3·93 (7·55) | 6·52 (4·93) | 6·83 (5·07) |
| **30-39 years** | 2·43 (7·49) | 3·33 (7·63) | 6·17 (4·87) | 6·64 (5·00) |
| **40-49 years** | 0·24 (7·63) | 1·34 (7·93) | 5·99 (4·72) | 6·54 (4·68) |
| **50-59 years** | -3·49 (7·87) | -3·75 (8·02) | 6·64 (5·47) | 7·11 (5·28) |
| **60-76 years** | -6·32 (7·65) | -6·47 (8·46) | 7·99 (5·87) | 8·47 (6·44) |

***Supplementary Table S6:* Mean absolute error (MAE) and brain predicted age difference (brain-PAD) per age group in the overall test samples of controls and major depressive disorder (MDD) patients.**
