## Supplementary_Appendix for "Brain Aging in Major Depressive Disorder: Results from the ENIGMA Major Depressive Disorder working group"

**Brain aging in Major Depressive Disorder - Supplementary appendix**

**Image exclusion criteria**

A neuroimaging expert at each scanning site inspected each image segmentation by overlaying the segmentation label of each structure on the T1-weighted brain scan. Additionally, study-wide statistics were collected (means and standard deviations) as well as histogram plots to identify non-normally distributed data and major outliers. Samples were excluded if its FreeSurfer feature was >2·698 standard deviations away from the global mean. If a sample was marked as a statistical outlier, the individual site was asked to re-inspect the subject’s segmentation in order to verify that it was properly segmented. If a sample was a statistical outlier, yet properly segmented, it was kept in the dataset. Otherwise, the sample was removed.

**Quality checking and sample exclusion criteria**

The initial dataset included 41 scanning sites with an age range of 7-89 years old based on N=8,728 samples (100%). However, due to scarcity of samples around the lower and upper age boundaries, we excluded those below 10 years (N=31) and above 75 years old (N=83), resulting in N=8,614 subjects (98·7%). Subsequently, the chronological age variable was floored, as some sites included one or two decimals in their age variable, while others did not. We checked individual FreeSurfer features for missings and excluded participant samples with >10% missing data, suggestive of poor reliability. This led to an exclusion of N=135 participants, resulting in the total sample of N=8,479 (97·1%).

The total sample of (N=8,479) was partitioned into datasets of controls and major depressive disorder (MDD) patients, separately for males and females, including N=2,301 (27·1%) male controls and N=1,265 (14·9%) male MDD patients, and N=2,745 (32·4%) female controls and N=2,168 (25·6%) female MDD patients. We divided healthy controls from each of the scanning sites into separate training (subset to train the model) and test samples (subset to test the trained model) using a balanced split-half approach. The 50:50 data partitioning was performed at random within each of these scanning sites while preserving the chronological age distribution between training and test data using the createDataPartition function from the “*caret*” package in R. Scanning sites with less than ten samples were excluded from the training sample, thus 28 scanning sites remained in the male sample (N=65 participants were excluded), compared to 34 scanning sites in the female sample (N=33 participants were excluded). The final training sample consisted of N=1,147 (13·5%) male controls and N=1,386 (16·3%) female controls. The final test samples consisted of N=1,089 (12·8%) male controls and N=1,167 (13·8%) depressed males and N=1,326 (15·6%) female controls and N=2,044 (24·1%) female depressed patients.

**Brain age prediction framework**

**Alternative machines/kernels**

To explore the effect of different machines and kernels, we repeated the 10-fold cross-validation training using gaussian process regression (GPR), ridge regression, and generalized additive models (GAM) in comparison to the support vector regression (SVR). To model non-linear multivariate patterns, we also explored radial basis function (RBF) kernels, as compared to linear kernel methods. Important to mention here is that all machine learning algorithms showed similar performances (**supplementary table S5**). Given the aim to make our model publicly available, we opted for the SVR with a linear kernel emphasizing its deployability and shareability. In contrast to the RBF kernels, linear kernels allow for sharing model weights at the feature level for making predictions in new independent test samples, without sharing any actual data points or support vectors from the training data. This ensures that no individual level data is shared.

**Associations between brain-PAD and FreeSurfer features**

Significantly associated FreeSurfer regions of interest (ROIs) with the brain-PAD outcome were plotted across groups (controls and MDD patients) in **figure 3** in the main manuscript, and separately in controls and MDD patients here (**supplementary figure S1**). There were no positive associations between brain-PAD and gray matter structures, with the exception of the mean lateral ventricles (controls: b=0·0004, p_FDR_=8·00^-35^; MDD: b=0·0005, p_FDR_=2·89^-60^). Intracranial volume (ICV) was also significantly negatively associated with brain-PAD in both controls and patients (controls: b=-3·45^-06^, p_FDR_=6·04^-06^; MDD: b=-5·34^-06^, p_FDR_=4·22^-14^), but was excluded from figure S1 for displaying purposes. We found strong widespread significant negative associations between brain-PAD and cortical thickness, and comparably weaker associations with surface area and subcortical volume measures in both groups. Important to note is the high degree of similarity of the strengths of the associations between brain-PAD and gray matter structures in controls and MDD patients.

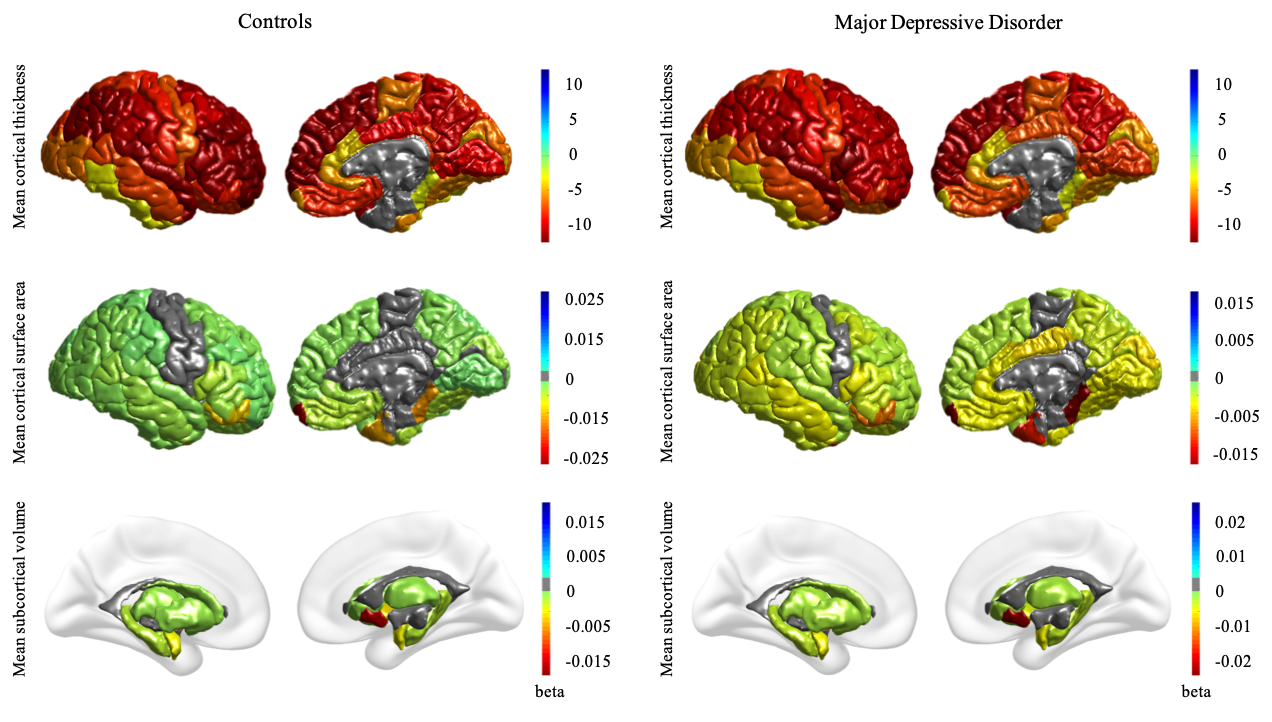

***Supplementary Figure S1:* Univariate associations between brain predicted age difference (predicted brain age - chronological age; brain-PAD) and FreeSurfer measures in controls (*left panel*) and major depressive disorder (MDD) patients (*right panel*).** Effect sizes (regression coefficient beta) are illustrated for regions with a significant (P_FDR_<0·05) negative association with brain-PAD, with the exception of a positive effect in the mean lateral ventricles. The figure shows associations with cortical thickness measures (*top row*), cortical surface areas (*middle row)*, and subcortical volumes (*bottom row)*. The brain-PAD estimates are adjusted for chronological age, age^2^, age^3^, sex and scanning site. The significant negative association with ICV was excluded from this figure for display purposes.

**Generalization to independent test samples from the ENIGMA Major Depressive Disorder working group**

The brain age prediction model generalized well to unseen samples (**supplementary table S6**). The overall correlations between predicted brain age and chronological age in the out-of-sample test controls from the ENIGMA MDD working group were *r*=0·87, *P*<0.001; R^2^=0·76 for males and r=0·86,p<0·001; R^2^=0·74 for females. Similarly, the performances in the MDD test samples were r=0·81, p<0·001; R^2^=0·66 for males, and r=0·82, p<0·001; R^2^=0·68) for females. Of note here is that prediction errors were not equal between sites and age groups (**supplementary figures S2-5**). More specifically, the mean absolute error (MAE) was highest in the oldest age group (60-75 years old, mean 8·28 [6·21]) and brain predicted age difference (brain-PAD) was significantly negatively associated with chronological age (overall r=-0·41, p<0.0001).

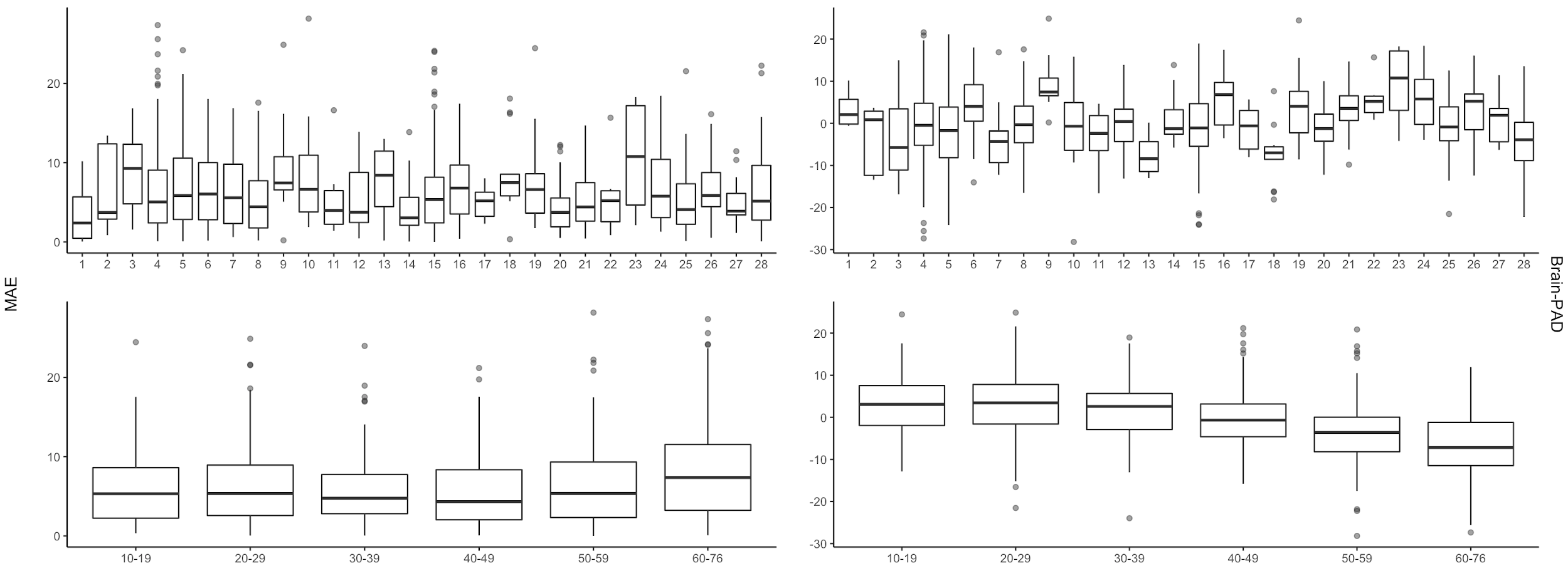

***Supplementary* *Figure S2*: Mean absolute error (MAE) and brain predicted age difference (brain-PAD) across scanning site and age group for the male control test samples.** Top row figures illustrate scanning sites on the x-axis. Prediction errors were examined across 28 different scanning sites and six different age groups of ten-year bins.

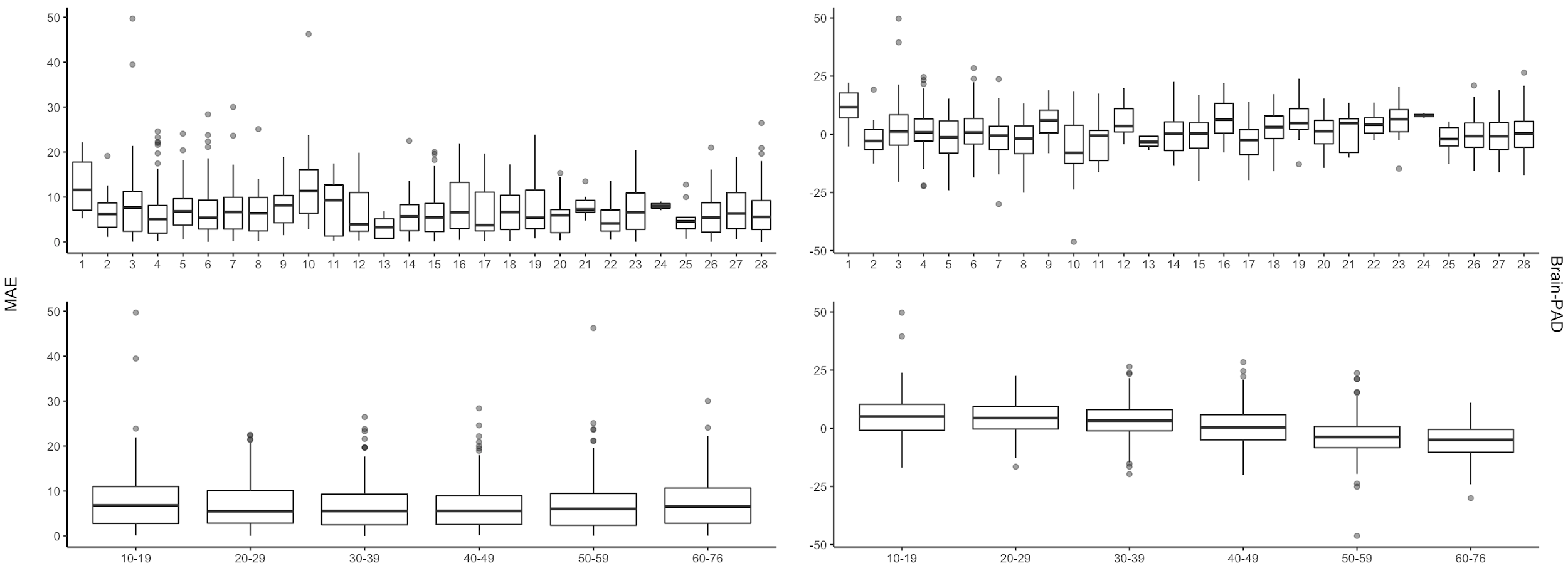

***Supplementary* *Figure S3*: Mean absolute error (MAE) and brain predicted age difference (brain-PAD) across scanning site and age group for the male major depression disorder (MDD) test samples.** Top row figures illustrate scanning sites on the x-axis. Prediction errors were examined across 28 different scanning sites and six different age groups of ten-year bins.

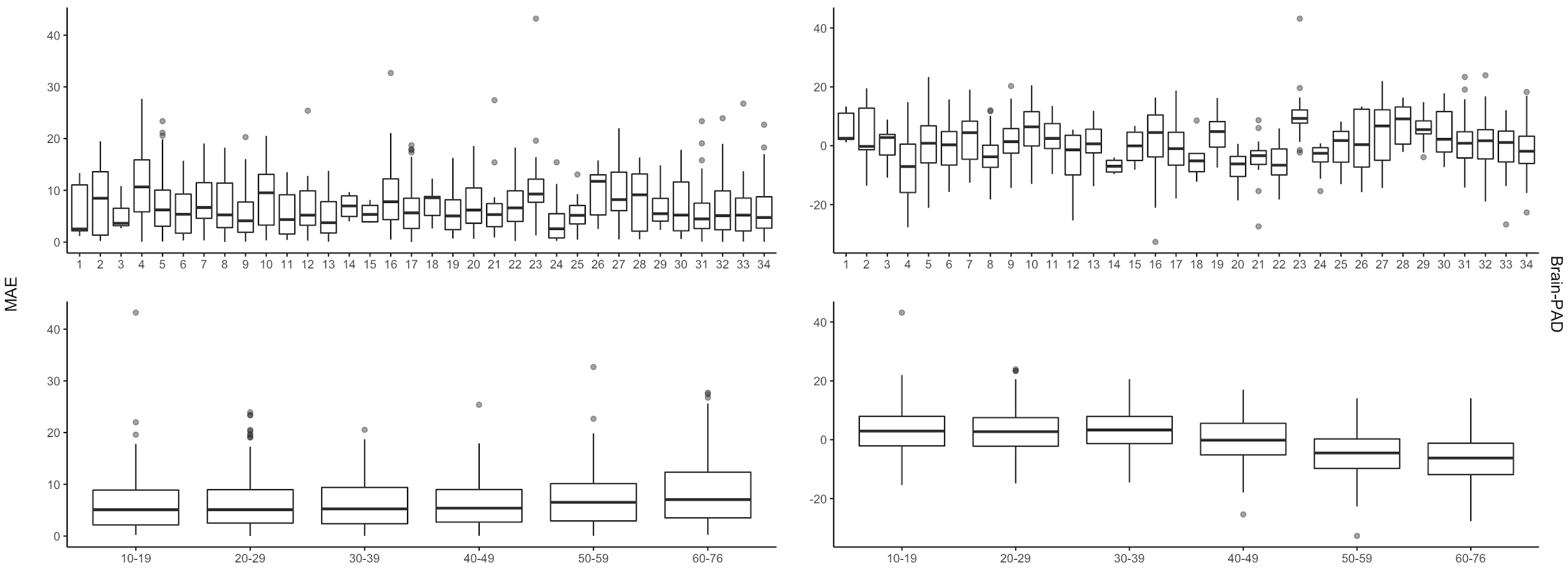

***Supplementary* *Figure S4*: Mean absolute error (MAE) and brain predicted age difference (brain-PAD) across scanning site and age group for the female control test samples.** Top row figures illustrate scanning sites on the x-axis. Prediction errors were examined across 34 different scanning sites and six different age groups of ten-year bins.

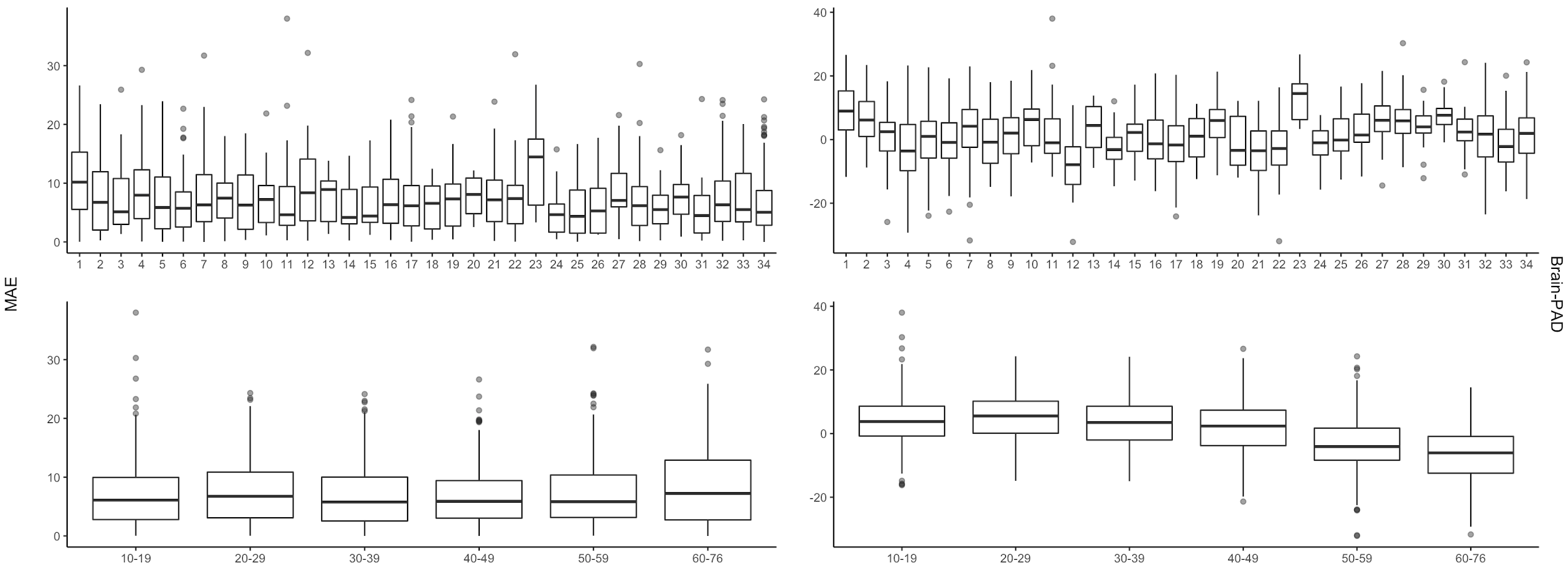

***Supplementary* *Figure S5*: Mean absolute error (MAE) and brain predicted age difference (brain-PAD) across scanning site and age group for the female major depression disorder (MDD) test samples.** Top row figures illustrate scanning sites on the x-axis. Prediction errors were examined across 34 different scanning sites and six different age groups of ten-year bins.

**Generalization to completely independent healthy controls from the ENIGMA Bipolar Disorder working group**

The brain age prediction models generalized well to healthy controls from completely independent samples (i.e. independent scanning sites) from the ENIGMA Bipolar Disorder (BD) working group (**supplementary figures 6-7**). The MAE was 7·24 (5·82) years in males and 7·45 (SD 5·44) in females, slightly higher than the MAE in the test samples of the ENIGMA MDD working group. The overall correlations between predicted brain age and chronological age in the out-of-sample controls were r=0·76, p<0.001; R^2^=0·57 for males and r=0·75, p<0·001; R^2^=0·56 for females (**supplementary figure S8**).

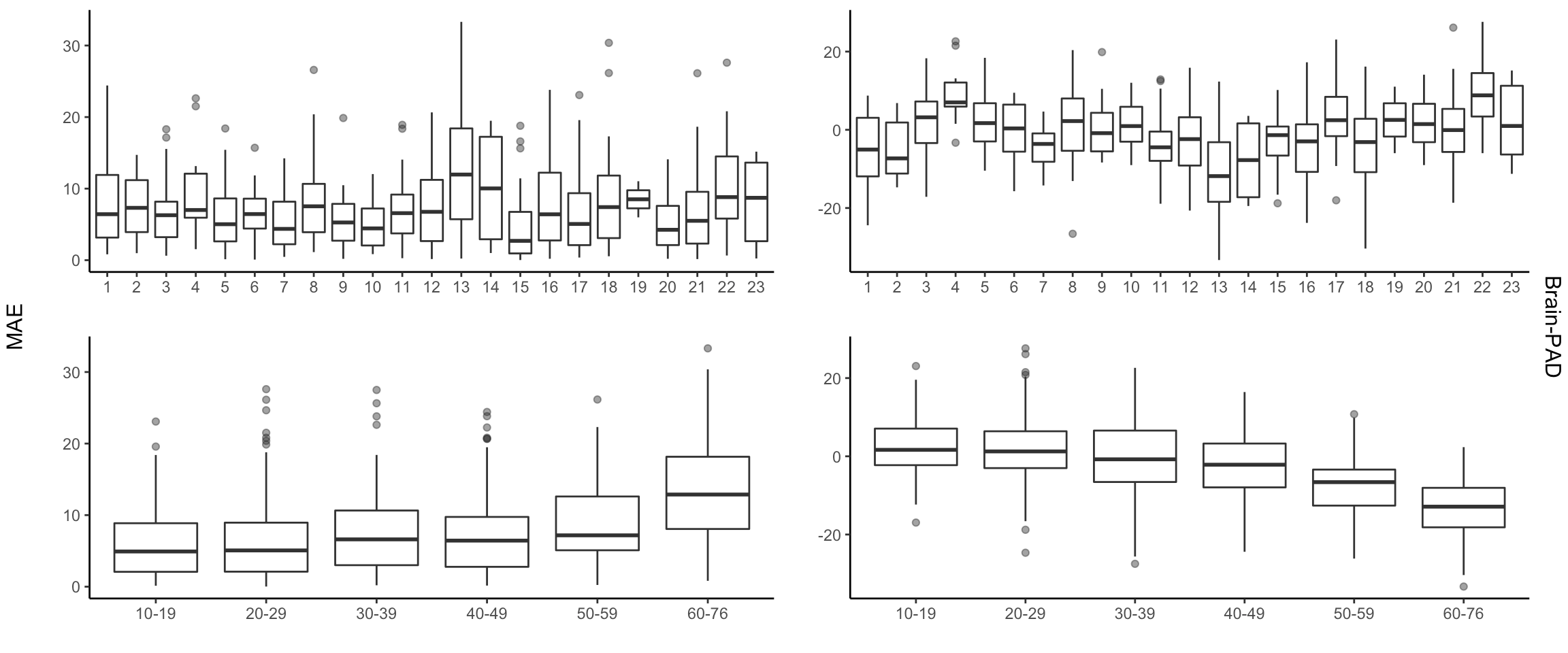

***Supplementary* *Figure S6*: Mean absolute error (MAE) and brain predicted age difference (brain-PAD) across scanning site and age group for the male control test sample from the ENIGMA Bipolar Disorder (BD) working group.** Top row figures illustrate scanning sites on the x-axis. Prediction errors were examined across 23 different scanning sites and six different age groups of ten-year bins.

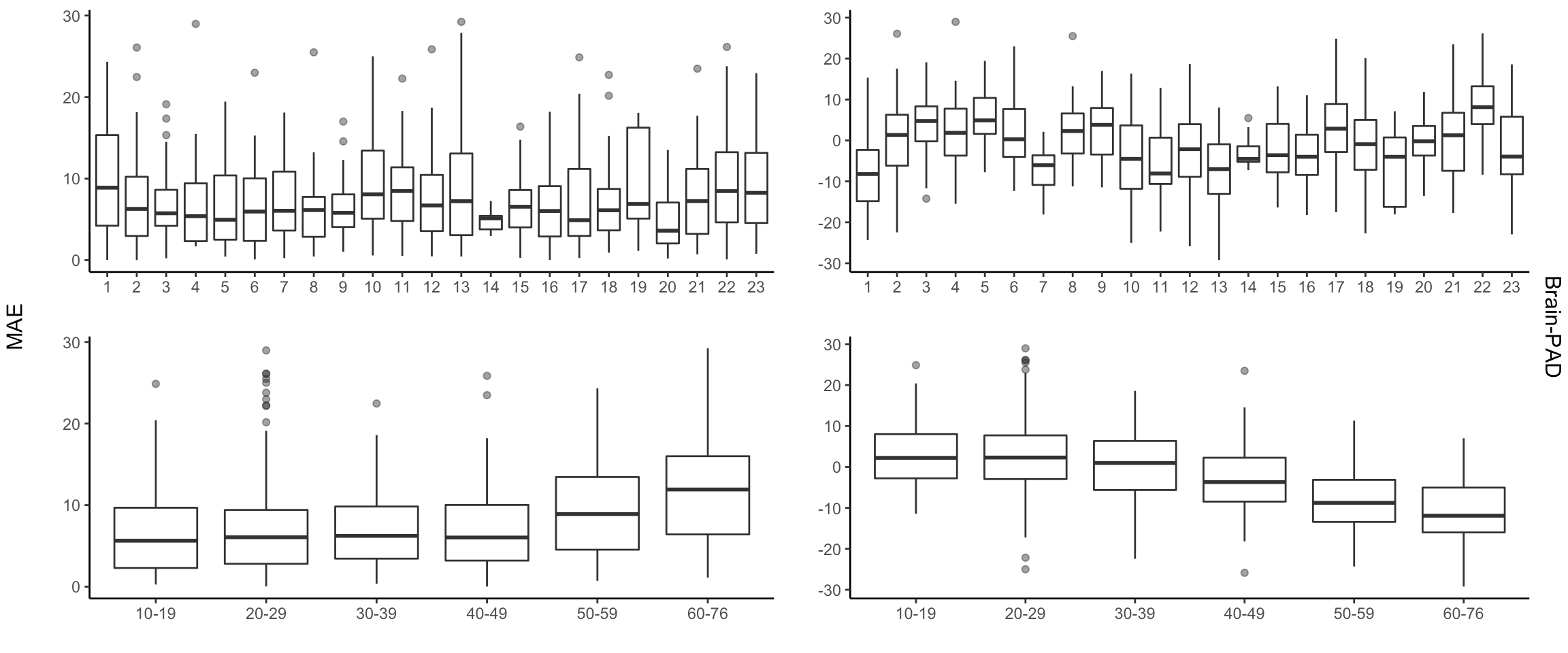

***Supplementary* *Figure S7*: Mean absolute error (MAE) and brain predicted age difference (brain-PAD) across scanning site and age group for the female control test sample from the ENIGMA Bipolar Disorder (BD) working group.** Top row figures illustrate scanning sites on the x-axis. Prediction errors were examined across 23 different scanning sites and six different age groups of ten-year bins.

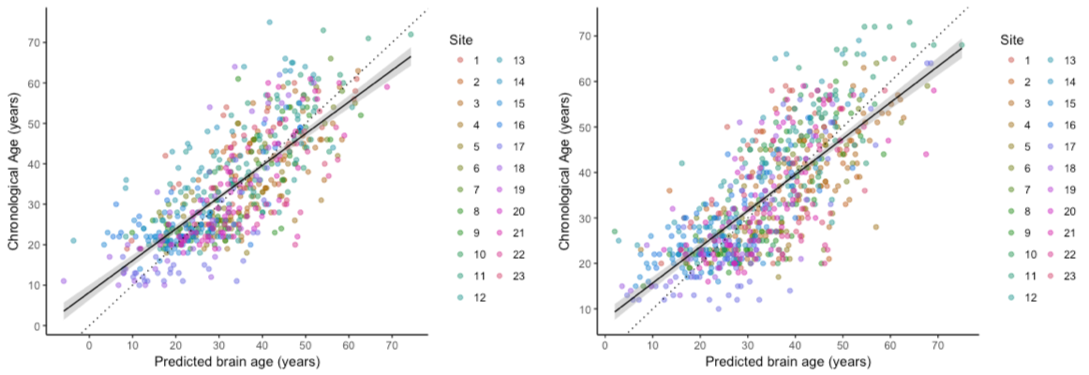

***Supplementary Figure S8:* Brain age prediction in completely independent control test samples from the ENIGMA Bipolar Disorder working group.** The plots show the correlation between chronological age and predicted brain age, derived from the 10-fold cross-validation of the Support Vector Regression model in the training samples of the ENIGMA MDD working group, separately for males (*left*) and females (*right*). The colors indicate scanning sites and each circle represents an individual subject. Diagonal dashed line reflects the line of identity (x=y).

**ENIGMA MDD Brain Age Model publicly available**

FreeSurfer is an automated and widely used software tool (<http://surfer.nmr.mgh.harvard.edu/>). Thus, our brain age algorithm can be easily applied to independent data, promoting validation and replication across different samples worldwide needed to mature modeling efforts, contributing to the development of canonical brain age models. To this aim, we will make our FreeSurfer-based brain age model publicly available at [https://www.photon-ai.com/](https://www.photon-ai.com/brainage) upon publication. Detailed instructions and guidelines for its use will be made available on the webpage. It is, however, important to note that prediction errors were higher in older age groups (>60 years old) and brain-PAD was significantly negatively associated with chronological age (r=-0·37 males, r=-0.40 females, both p’s<0.0001), with the latter being a known feature of the brain-PAD metric.^1^ Thus, caution is warranted when applying our model to data from older participants (>60 years). We recommend to: a) only use our models to samples with an upper age limit of 60 years, and b) always include residual chronological age effects as covariates in the analyses.

**References**

1	Le TT, Kuplicki RT, McKinney BA, *et al.* A Nonlinear Simulation Framework Supports Adjusting for Age When Analyzing BrainAGE. *Front Aging Neurosci* 2018; **10**: 317.
